# Subgraphs of functional brain networks identify dynamical constraints of cognitive control

**DOI:** 10.1101/147272

**Authors:** Ankit N. Khambhati, John D. Medaglia, Elisabeth A. Karuza, Sharon L. Thompson-Schill, Danielle S. Bassett

## Abstract

Brain anatomy and physiology support the human ability to navigate a complex space of perceptions and actions. To maneuver across an ever-changing landscape of mental states, the brain invokes cognitive control – a set of dynamic processes that engage and disengage different sets of brain regions to modulate attention, switch between tasks, and inhibit prepotent responses. Current theory suggests that cooperative and competitive interactions between brain areas may mediate processes of network reorganization that support transitions between dynamical states. In this study, we used a quantitative approach to identify distinct topological states of functional interactions and examine how their expression relates to cognitive control processes and behavior. In particular, we acquired fMRI BOLD signal in twenty–eight healthy subjects as they performed two cognitive control tasks – a local-global perception switching task using Navon figures and a Stroop interference task – each with low cognitive control demand and high cognitive control demand conditions. Based on these data, we constructed dynamic functional brain networks and used a parts-based network decomposition technique called non-negative matrix factorization to identify putative cognitive control subgraphs whose temporal expression captured key dynamical states involved in control processes. Our results demonstrate that the temporal expression of these functional subgraphs reflect cognitive demands and are associated with individual differences in task-based performance. These findings offer insight into how coordinated changes in the cooperative and competitive roles of distributed brain networks map trajectories between cognitively demanding brain states.

## 1. Introduction

In human cognition, internally-generated *cognitive control* processes modulate attention, facilitate task switching, and inhibit prepotent behavior (Medaglia et al., 2016a). One avenue by which the brain may rapidly traverse a cognitive state-space is through its functional interactions – coherent fluctuations in brain activity mediated by the structural connectome (Deco et al., 2011). The brain’s distributed functional interactions form a functional network whose architectural configuration is temporally dynamic (Hutchison et al., 2013), conferring flexible adaptivity in the face of environmental pressures or task demands (Bassett et al., 2006) such as those elicited during learning (Bassett et al., 2011) and other tasks demanding executive cognition (Braun et al., 2015). Cognitive control processes have been widely reputed to recruit several cognitive systems that include executive, attention, and salience systems that span prefrontal cortices, striatum, parietal regions and cerebellum (Duncan and Owen, 2000; Miller, 2000; Koechlin et al., 2003; Liston et al., 2006; Ito, 2008; Krienen and Buckner, 2009). The notion that cognitive control involves a heterogenous collection of brain systems is robustly supported by several univariate studies that robustly demonstrate concurrent activation of functionally-specialized brain areas across different cognitive control tasks (Cole and Schneider, 2007). If similar sets of brain areas appear to be active across a diverse set of cognitive control tasks, then how do functional brain networks encode nuanced differences between tasks and adapt to fluctuations in cognitive demand (Fig. 1A)?

**Figure 1:**
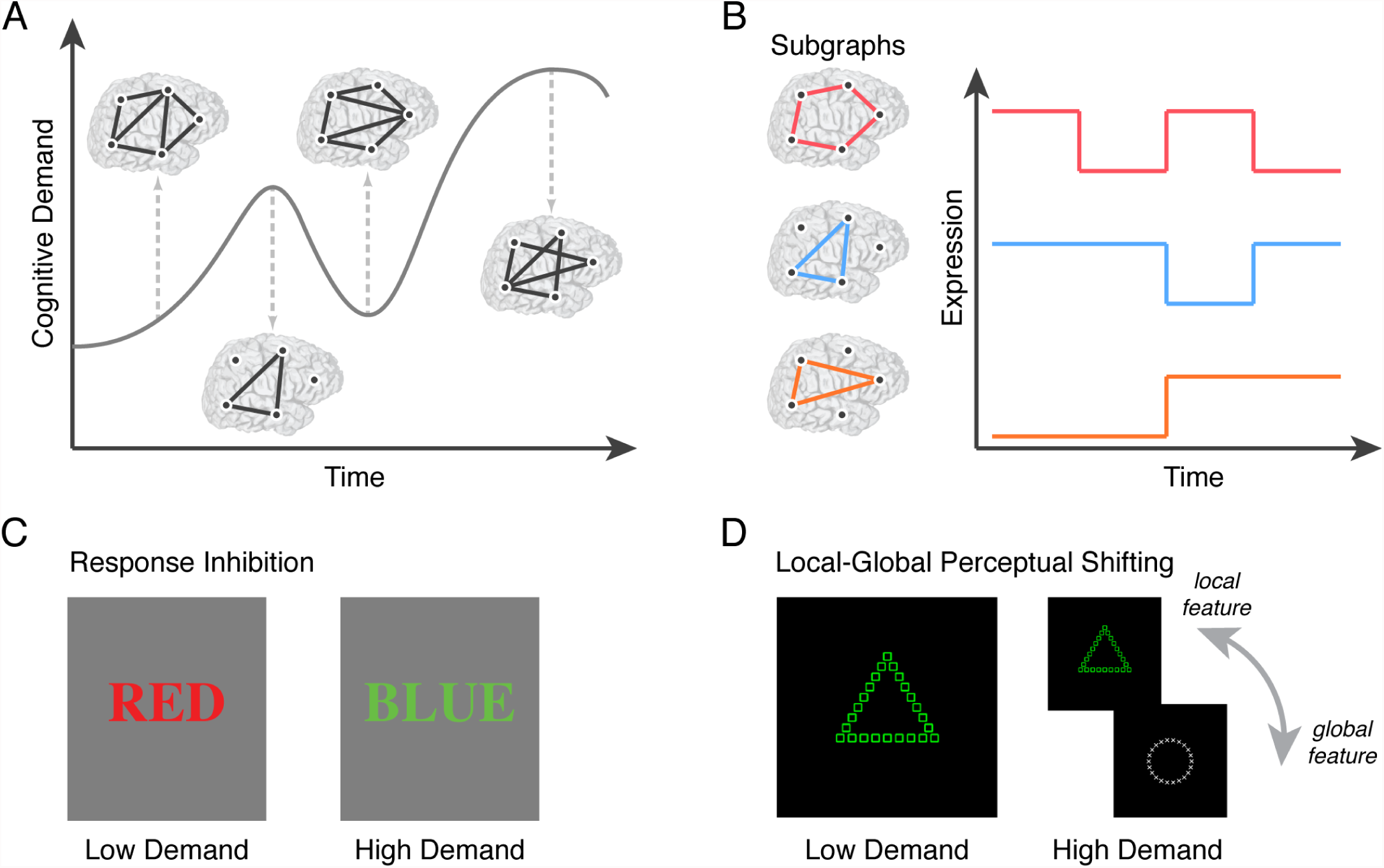
Experimentally modulating cognitive control processes to uncover internal mechanisms of network regulation. (*A*) To monitor and regulate the demands placed on neural systems, empirical evidence suggests that the brain employs putative *cognitive control* processes that gate information and select among competing representations and processes (Botvinick et al., 2001). Functional brain networks that flexibly coordinate interactions between different sets of brain regions over time may be a key substrate for cognitive control, and moreover be essential for maintaining homeostasis between internally-driven brain dynamics and externally-elicited behavioral goals (Miller, 2000). A graph theoretical framework helps us model the dynamics of cognitive control networks by representing brain regions as *nodes* and the strength of functional interactions between brain regions as *weighted edges*. (*B*) Recent advances in network neuroscience (Bassett and Sporns, 2017) and machine learning (Khambhati et al., 2016b) enable us to cluster functional brain networks into composite subgraphs – cohesive sets of graph edges (*left*) from the observed network (*A*) that tend to co-vary in strength over time. The putative role of a subgraph in cognitive control is inferred by its relative level of weighted expression in the observed network at a specific task block during cognitive processing (*right*). To experimentally modulate cognitive demand, we recruited 28 healthy adult human participants to perform a response inhibition, Stroop task (*C*) and a task-switching, local-global feature perception task based on Navon figures (*D*). The Stroop task entailed (i) a fixation condition consisting of a black crosshair at the center of the screen, (ii) a low demand condition consisting of a matched word-color pair, and (iii) a high demand, interference condition consisting of a mismatched word-color pair. Subjects were required to report the color of the presented word. The Navon task entailed (i) a fixation condition consisting of a black crosshair at the center of the screen, (ii) a low demand condition consisting of only white or green Navon figures – local shapes embedded in a non-matching global shape, and (iii) a high demand condition consisting of Navon figures randomly alternating between white or green color. Subjects were required to report the local shape if the presented figure was white or to report the global shape if the presented figure was green. Differences in block types in each control task are thought to invoke different levels of recruitment of cognitive control mechanisms. Participant reaction time on correct trials was used to measure performance, and the difference in performance between high and low cognitive control blocks is thought to represent the costs of cognitive control.

One mechanistic theory, known as the “adaptive coding model of cognitive control” (Duncan, 2001), posits that brain regions that activate during higher cognitive functions can alter their dynamical properties based on the current goals of the neural system. More recent studies have challenged this hypothesis by presenting data that suggests that changes in the cognitive demands of a task lead to recruitment of mechanistically-specialized brain regions based on an anatomically-defined gradient (Badre and D’Esposito, 2007, 2009). To reconcile these opposing theories of the neuronal basis of cognitive control, Woolgar et al. (2015) applied multivoxel pattern analysis – a machine learning technique for identifying consistent patterns of voxel-wise activation – to the fMRI of subjects as they performed simple and cognitively demanding tasks. The authors found a consistent pattern of activation in frontoparietal brain areas that was specific to highly demanding conditions across multiple cognitive tasks. Their findings support the hypothesis that a consistent group of brain regions activate in response to increases in cognitive demand. However, parallel lines of investigation on the underpinnings of cognitive control in functional brain networks suggest that the integrated cognitive control network dissociates into several, segregated sub-networks that are responsible for different aspects of cognitive control processes (Cole and Schneider, 2007). To address these conflicting reports, a data-driven approach that can disentangle functional interactions that encode cognitive states associated with control tasks and track their expression alongside changes in cognitive demand is required. Such a capability would improve our understanding of which components of functional brain networks are important for different facets of cognitive control, and how these components encode shifts between cognitively demanding states.

In the present work, we identified components of functional brain networks that facilitate the transition between cognitively demanding states by using an unsupervised machine learning technique known as non-negative matrix factorization (NMF) (Lee et al., 1999). NMF decomposes functional brain networks into: (i) additive subgraphs that represent clusters of graph edges that track with one another over time, and (ii) time-varying coefficients that quantify the degree to which a subgraph is expressed at a point in time (Khambhati et al., 2016b; Chai et al., 2017; Khambhati et al., 2017). This computational tool allowed us to trace how groups of functionally interacting brain areas are dynamically expressed during experimentally modulated changes in cognitive demand (Fig. 1B). In particular, participants engaged in the following two cognitive control tasks: a response inhibition Stroop task (Fig. 1C; Stroop (1935)) and a local-global perception switching task based on classical Navon figures (Fig. 1D; Navon (1977)). Our methodological approach enabled us to address a critical question in cognitive control: “How do brain networks adapt to the cognitive demands imposed by a particular task?”

To address this question using NMF, we drew upon recent studies that suggest task-driven reconfiguration of functional brain networks integrates otherwise functionally-specialized and segregated information (Shine et al., 2016; Bassett et al., 2015). One compelling current theory posits that transitions between cognitively demanding brain states may be mediated by dynamic changes in the patterns of competitive and cooperative functional interactions: that is, between (i) anticorrelated fluctuations in activity between brain regions that represent segregated brain functions, and (ii) correlated fluctuations in activity between brain regions that represent integrated brain functions (Fornito et al., 2012; Cocchi et al., 2013). Competitive and cooperative dynamics are putative mechanisms for how task-relevant information is transferred between different regions of the network during cognitively demanding tasks. By accounting for the cooperative or competitive nature of functional interactions in the NMF framework, we were able to distinguish the likelihood that the functional interactions within a subgraph were competitively or cooperatively expressed at a particular point in time – providing a perspective on integrated and segregated information processing of composite sets of brain regions.

Based on prior studies demonstrating that behavioral tasks can be used to dissociate intrinsic and task-specific architectures of functional brain networks (Cole et al., 2014), we first hypothesized that NMF would identify functional subgraphs whose expression is either generalized across the Stroop and Navon tasks or specific to distinct cognitive conditions within and between tasks. In particular, we expected task-general subgraphs to reflect interactions relevant for task saliency and cognitive control processes common to both tasks; we also expected task-specific subgraphs to reflect interactions relevant for stimulus processing and attentional mechanisms necessary for either response inhibition in the Stroop task or task-switching in the Navon task. Building upon recent evidence that functional interactions dynamically reorganize between co-operatively integrated and competitively segregated brain states (Shine et al., 2016), we next hypothesized that functional subgraphs would shift their roles between cooperative and competitive modes of interaction in response to experimentally driven changes in cognitive demand. Lastly, we hypothesized that changes in subgraphs expression during experimental modulation of cognitive demand would discriminate inter-individual differences in behavioral performance on the task. Specifically, based on previous theories regarding the behavioral influence of cooperative and competitive functional interactions in cognitive control (Cocchi et al., 2013), we expected that components of the executive system would most prominently participate in subgraphs associated with individual differences in performance.

## 2. Results

### 2.1. Decomposing functional subgraphs of cognitive control

To uncover the putative roles of cooperative and competitive functional interactions in cognitive control, we first performed fMRI while 30 healthy adult human subjects performed Stroop and Navon cognitive control tasks. Two subjects were excluded on the basis of poor performance and technical problems on the day of scanning, leaving 28 subjects for further analysis. In particular, we measured fMRI BOLD signals from 262 functional brain areas (Fig. 2A) – including cortex, subcortex, and cerebellum (Cammoun et al., 2012; Diedrichsen et al., 2009) – during three separate conditions of both the Stroop and Navon tasks: fixation, low cognitive demand, and high cognitive demand conditions (Fig. 2B). Briefly, the low cognitive demand condition was designed to elicit a neural response associated with performing each task with low cognitive control demands and the high cognitive demand condition was designed to elicit a neural response associated with either task shifting or inhibition cost (see *Experimental Procedures* for more details). We then constructed dynamic functional brain networks for each subject where network nodes represent brain regions and network edges between nodes represent the Pearson correlation coefficient between regional BOLD time series (Fig. 2C). Specifically, we computed a 262 × 262 adjacency matrix for each of 6 task blocks (corresponding to 30 seconds of BOLD activity, and comprising several trials) in each of the 3 task conditions (fixation, low demand, high demand) for each of 2 tasks (Stroop and Navon). This process results in 36 block-level adjacency matrices per subject. Importantly, positive Pearson correlations underlie integrated and coherent activation between brain regions or *cooperative functional interactions* and negative Pearson correlations underlie segregated and discordant activation between brain regions or *competitive functional interactions* (Fornito et al., 2012) (see Fig. S1 for the distribution of correlations between brain regions). To separate positively-weighted network edges (cooperative interactions) from negatively-weighted network edges (competitive interactions), we duplicated the adjacency matrix of each block and separately thresholded edge weights either greater than zero or less than zero (see *Experimental Procedures* for more details). Lastly, we aggregated all functional brain networks into a network configuration matrix (Fig. 2D) with size 2016 × 34191. The first dimension of size 2016 corresponds to all combinations of two tasks, three task conditions, six repeated blocks, twenty-eight subjects, and two edge types (cooperative or competitive); the second dimension of size 34191 corresponds to all unique, pairwise edges between the 262 brain regions.

**Figure 2:**
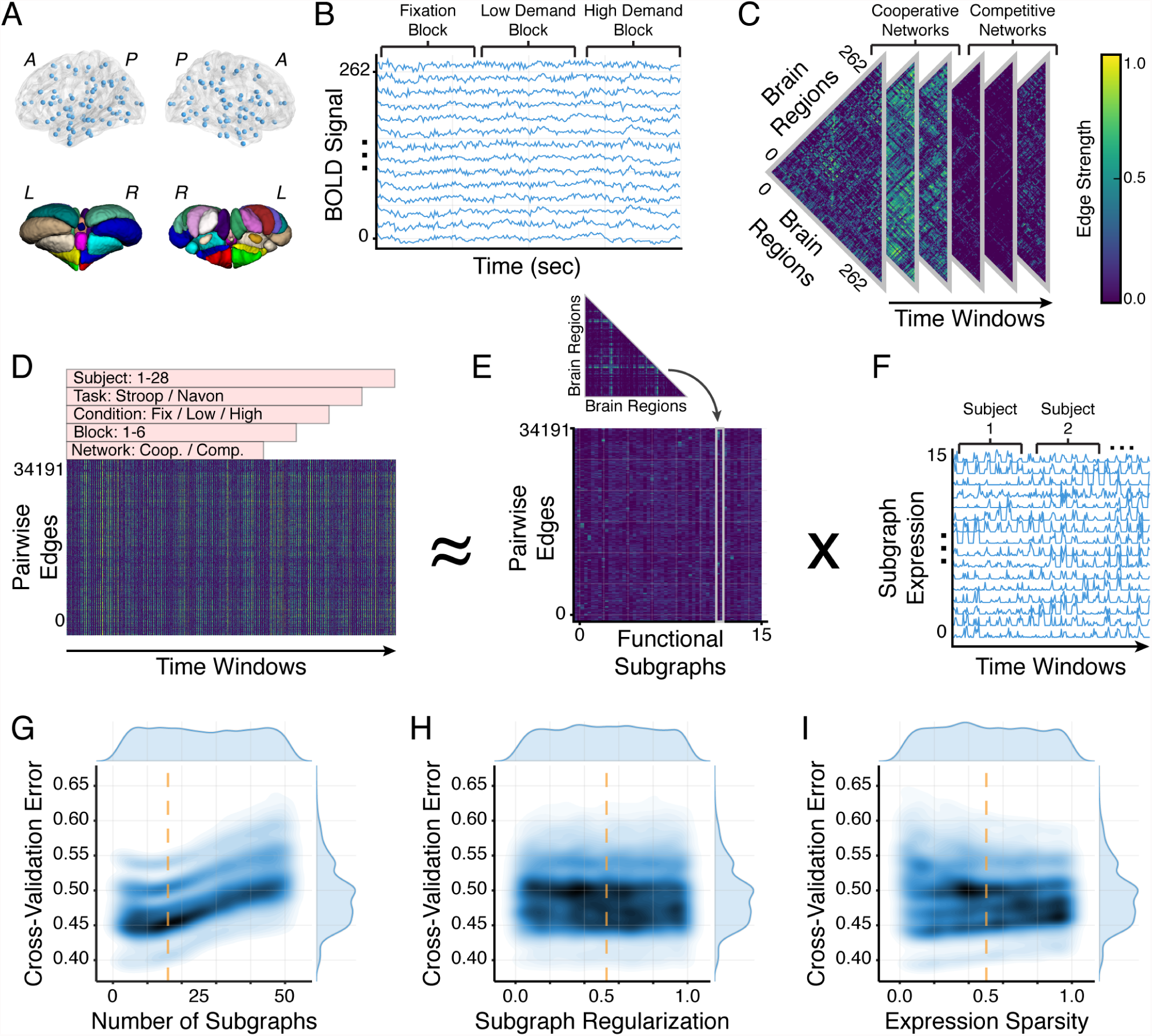
Learning functional subgraph architecture of cognitive control processes. (*A*) We obtained fMRI BOLD signals from 262 functional regions of interest (234 cortical and subcortical brain areas parcellated by (Cammoun et al. (2012); *top*) and 28 cerebellar brain areas parcellated by (Diedrichsen et al. (2009); *bottom*) as 28 healthy adult human subjects performed Stroop and Navon cognitive control tasks. (*B*) We concatenated BOLD signal from 6 task blocks (corresponding to 30 seconds of BOLD activity, and comprising several trials) in each of 3 task conditions (fixation, low demand, high demand) for each of 2 tasks (Stroop and Navon). (*C*) Next, we calculated the Pearson correlation coefficient between each pair of regional BOLD signals to create an adjacency matrix for every experimental block. We encoded this information in dynamic functional networks with brain regions as graph nodes and time-varying coherence as weighted graph edges. To assess the relative impact of *cooperative* interactions (positively weighted edges) and *competitive* interactions (negatively weighted edges) on cognitive control, we thresholded each adjacency matrix and separately grouped positive edges and negative edges (see *Experimental Procedures*). (*D*) We concatenated all pairwise edges over task blocks and subjects, and we generated a single time-varying network configuration matrix for the entire study cohort (*right*). We applied non-negative matrix factorization (NMF) – which pursues a parts-based decomposition of the dynamic network – to the configuration matrix and clustered temporally co-varying graph edges into a matrix of subgraphs (*E*) and a matrix of time-dependent coefficients (*F*) that quantify the level of expression in each task block for each subgraph. (*G-I*) NMF-based subgraph detection requires optimizing three parameters: the number of subgraphs *m*, the temporal sparsity of subgraph expression *β*, and the regularization of subgraph edge weights *α*. To characterize this parameter space, we randomly sampled *m*, *β*, and *α* from a three-dimensional uniform distribution (*m* [2, 50], *β* ∊ [0.01, 1.0], *α* ∊ [0.01, 1.0]) and applied NMF to the configuration matrix using each parameter set. Kernel density estimate of each bivariate distribution is indicated by the contour plot, where darker shades of blue indicate greater probability mass of the random sampling distribution. Optimal parameters are the average parameter values that yielded cross-validation error in the bottom 25% of the sampling distribution and are indicated by the dashed orange line.

To disentangle functional subgraphs and their dynamic expression from functional brain networks, we applied an unsupervised machine learning algorithm called non-negative matrix factorization (NMF) to the network configuration matrix. This technique enabled us to pursue a parts-based decomposition of network edges into additive functional subgraphs (Fig. 2E) with accompanying expression coefficients that measure the degree to which the subgraph is expressed in a particular task block, task condition, subject, and edge type (Fig. 2F) (Chai et al., 2017). Each subgraph composes a 262 × 262 adjacency matrix and each subgraph’s expression coefficients compose a vector of length 2016. Thus, subgraphs detail topological components of the functional brain network and the temporal coefficients quantify their expression during different phases of the cognitive control tasks. Moreover, each subgraph was associated with a cooperative expression component (based on positively-weighted network edges) and a competitive expression component (based on negatively-weighted network edges). A critical step in using NMF is optimizing model parameters (number of subgraphs *m*, temporal sparsity of subgraph expression *β*, and regularization of subgraph edge weights *α*) to ensure generalizability of component subgraphs without overfitting the model on observed data. By designing a fourteen-fold, leave-two-subjects-out cross-validation scheme, we minimized the average cross-validation error on held-out subjects and found the optimal number of subgraphs to be sixteen, the temporal sparsity to be 0.50, and the regularization of the subgraph edge weights to be 0.52 (Fig. 2G-I; see *Experimental Procedures* for more details). Finally, we ranked subgraphs (*A-P*) in increasing order of their generalized expression across all conditions in the cognitive control tasks. Specifically, we computed the expression sparsity for each subgraph as its mean number of expression coefficients with zero weight (Fig. S2). Intuitively, subgraphs with greater expression sparsity are more specific to particular conditions of the cognitive control tasks while subgraphs with lower expression sparsity are less specific to any particular task condition and more generalized to the full set of tasks. We referred to specific subgraphs according to their assigned letter for the remainder of the study.

We first asked whether the functional subgraphs expressed during the cognitive control tasks reflected functional interactions within and across known cognitive systems. To study the relationship between the functional subgraph architecture and known cognitive brain systems, we assigned each of the 262 brain regions into one of eleven cognitive systems (Gu et al., 2015): frontoparietal, cingulo-opercular, dorsal attention, ventral attention, default mode, somatosensory, auditory, visual, subcortical, cerebellum, and other (see Table S1 for specific region-to-system assignments). Thus, we re-organized the rows and columns of each subgraph’s 262 × 262 adjacency matrix such that nodes assigned to the same brain system were contiguously ordered, and we then visualized the resulting adjacency matrices as circular, ring graphs (Fig. 3). To quantitatively confirm that each subgraph captured functional interactions that were indeed distributed within and between cognitive systems, we compared the average subgraph edge weight between pairs of nodes of the same or different cognitive system to a null distribution of the average subgraph edge weight – constructed by permuting subgraph edge weights between nodes and recomputing the average subgraph edge weight for each pair of cognitive systems for 1000 permutations. We found that functional subgraphs clustered interactions between brain regions of the same cognitive system and between brain regions of different cognitive systems (*p* < 0.05; Bonferroni corrected for multiple comparisons; see Fig. S3) – implicating a distributed functional architecture underlying the cognitive control tasks. In other words, the functional subgraphs recovered by NMF span several cognitive brain systems defined *a priori* (Gu et al., 2015).

**Figure 3:**
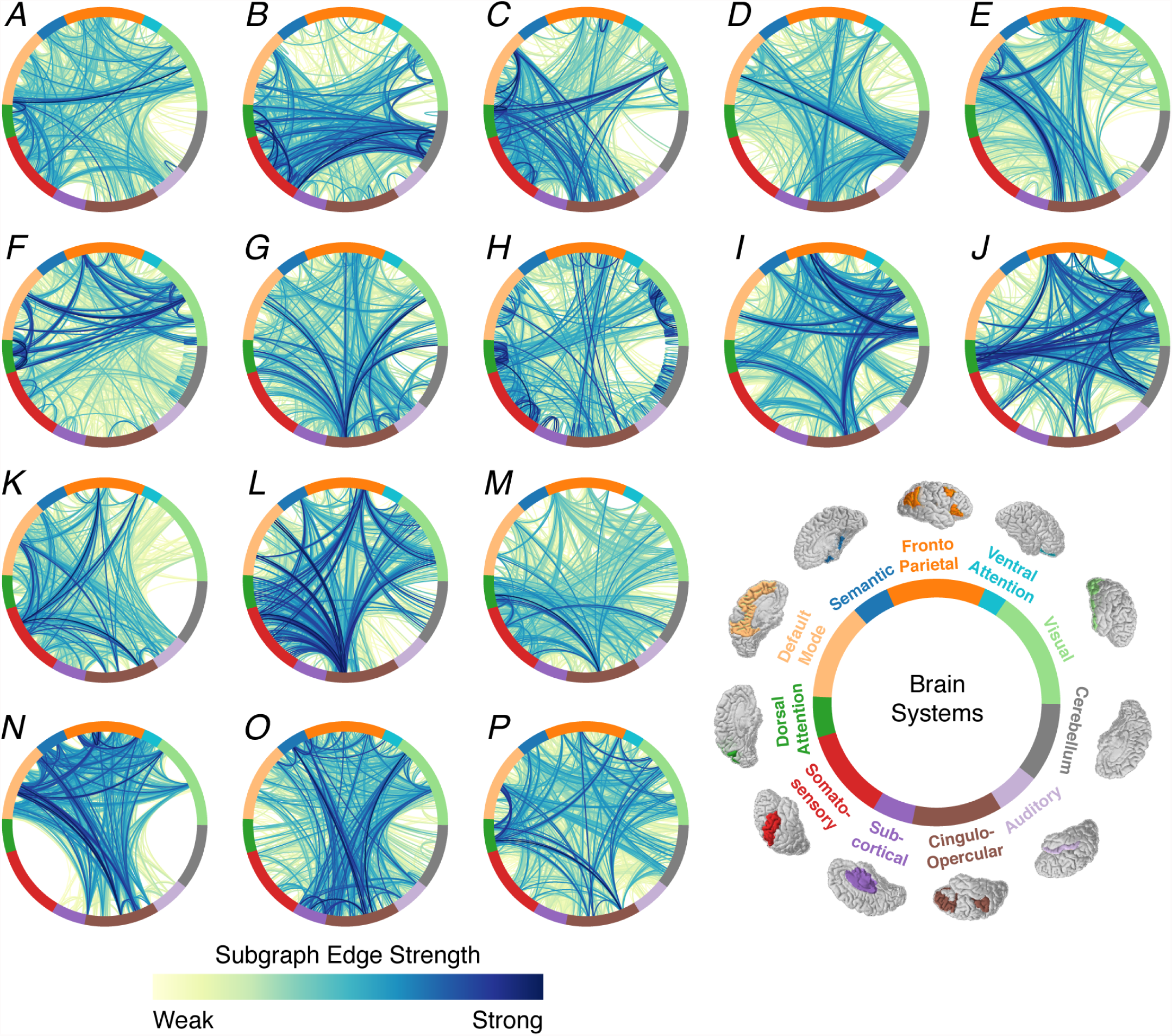
Linking functional subgraphs to neuroanatomy of canonical cognitive systems. By applying NMF to the network configuration matrix, we uncovered sixteen functional subgraphs whose weighted graph edges span between the 262 graph nodes specified by the brain atlas. To examine functional roles of subgraphs in cognitive processing, we assigned each node in the atlas to one of the following eleven putative cognitive systems (Gu et al., 2015): frontoparietal, cingulo-opercular, dorsal attention, ventral attention, default mode, somatosensory, auditory, visual, subcortical, cerebellum, and other. To visualize the topological architecture of each subgraph, we constructed ring graphs in which nodes are evenly spaced around a circle – ordered and color coded by assigned cognitive system - and edges between nodes are represented by line arcs – colored by the percentile of the edge strength in the subgraph. Subgraphs coded *A* through *P* in increasing order of the percentage of sparse temporal expression coefficients – subgraphs closer to *A* tend to be expressed more generally across task conditions and subgraphs closer to *P* tend to be expressed more specifically in particular task conditions. For system-by-system adjacency matrix representations of functional subgraphs, we refer the reader to Fig. S3.

### 2.2. Recruitment of functional subgraphs during cognitive control tasks

Based on the set of sixteen functional subgraphs and their temporal expression, we next asked “Are functional subgraphs differentially recruited during separate cognitive control tasks?” We hypothesized that a functional subgraph would be sensitive to either cognitive processes specific to each task or cognitive control mechanisms that overlap between the tasks. To test our hypothesis, we compared relative differences of each subgraph’s expression between the Stroop task and the Navon task. Specifically, we computed the average subgraph expression over all blocks and conditions of each task for every subject and subtracted the average subgraph expression for the Navon task from the average subgraph expression on the Stroop task for every subject. Negative values of this difference suggest greater subgraph expression during the Navon task and positive values of this difference suggest greater subgraph expression during the Stroop task. We separately computed the relative subgraph expression for cooperative expression coefficients and competitive expression coefficients. We then ranked the subgraphs in order of increasing mean relative subgraph expression over all subjects (Fig. 4). Using a one-way ANOVA, we assessed whether subgraphs are expressed differently depending on the cognitive control task. We indeed found that subgraphs generally differ in their cooperative expression (*F*_15_ = 4.7, *p* = 2.0 × 10^−8^) and their competitive expression (*F*_15_ = 4.5, *p* = 4.9×10^−8^)acrosstheStroopandNavontasks. Theseresults suggest that functional subgraphs differentiate clusters of functional interactions that are separately relevant for the Stroop or Navon cognitive control tasks. This also implies that the two cognitive control tasks are uniquely stereotyped by clusters of cooperative functional interactions and competitive functional interactions.

**Figure 4:**
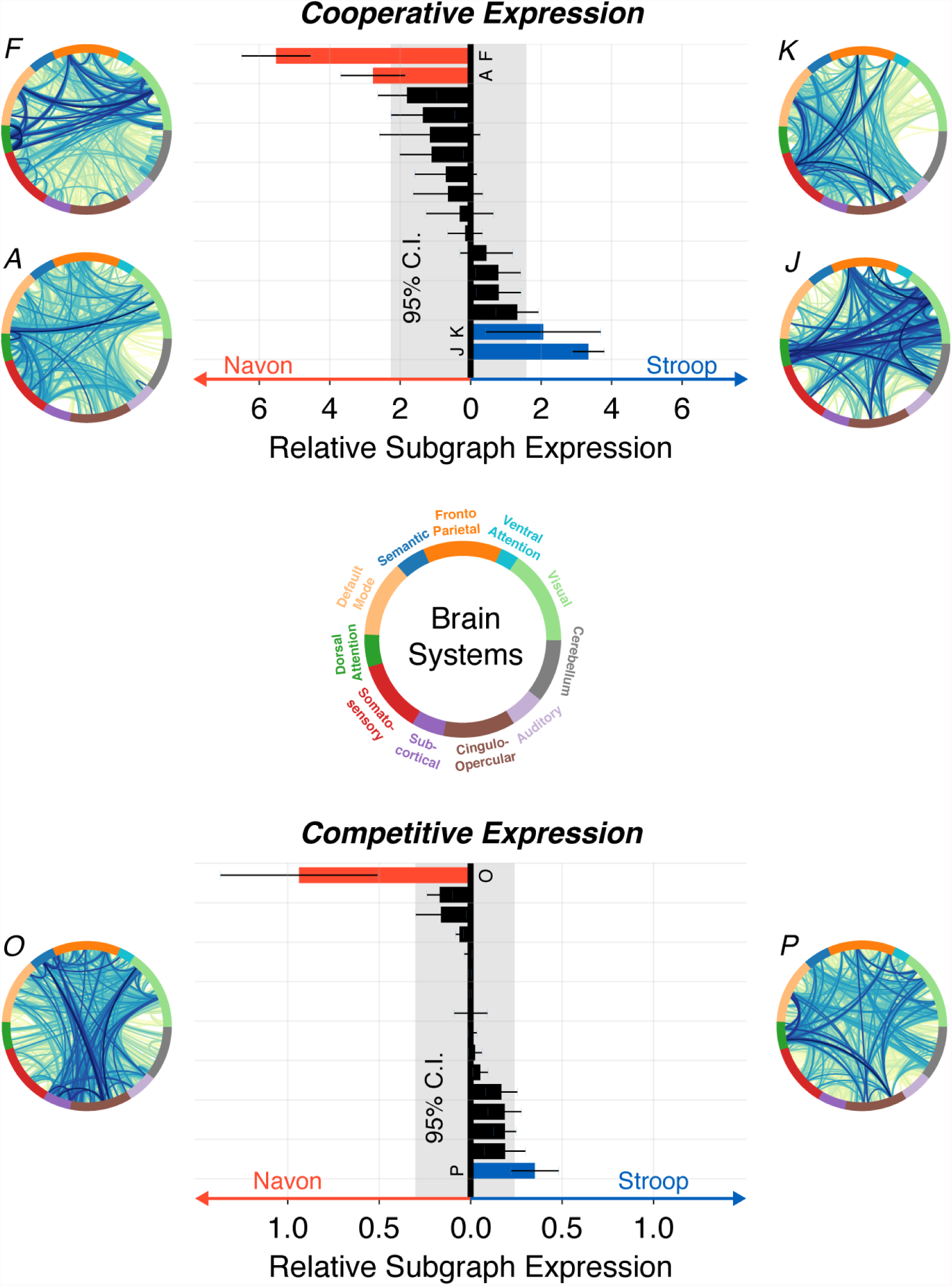
Subgraphs map functional interactions specific to cognitive control tasks. Distributions of the relative subgraph expression between Navon and Stroop tasks for cooperative functional interactions (*top*) and competitive functional interactions (*bottom*). Each bar represents the difference of a subgraph’s mean expression – averaged over all task conditions and blocks - between the Navon task and Stroop task, averaged over all subjects. Subgraphs with negative expression difference exhibited more expression bias for the Navon task and subgraphs with positive expression difference exhibited more expression bias for the Stroop task. Using a one-way ANOVA test, we found that subgraphs are differentially expressed during the Stroop and Navon tasks for cooperative interactions (*F*_15_ = 4.7, *p* = 2.0×10^−8^) and for competitive interactions (*F*_15_ = 4.5, *p* = 4.9×10^−8^). To determine whether a subgraph was more expressed for a task than would be expected by chance, we compared the relative subgraph expression to a null distribution in which task-specific differences in subgraph expression were uniformly permuted across subgraph assignments and subjects, and averaged over all subjects, 1000 times. For the Navon task, we found that cooperative interactions of subgraphs *F* and *A* and competitive interactions of subgraph *O* expressed significantly greater task-specific bias than expected by chance (*p* < 0.025). For the Stroop task, we found that cooperative interactions of subgraphs *K* and *J* and competitive interactions of subgraph *P* expressed significantly greater task-specific bias than expected by chance (*p* < 0.025).

We next asked which functional subgraphs were most significantly biased for expression during either the Stroop task or the Navon task. To determine whether any particular subgraph was more expressed for one of the tasks than expected by chance, we generated a null distribution of relative subgraph expression using a permutation test in which we uniformly redistributed subgraph expression coefficients across all tasks, task conditions, and subjects, and then recomputed the relative subgraph expression for each of 1000 permutations. We found three subgraphs (*F, A, O*) with significantly greater expression bias for the Navon task than expected by chance (*p* < 0.025; Bonferroni corrected for multiple comparisons). Specifically, subgraphs with significantly biased expression towards the Navon task conditions exhibited: (i) cooperative interactions between visual, fronto-parietal, cingulo-opercular, and default mode systems (subgraph *F*), (ii) cooperative interactions between visual, somatosensory, auditory, and default mode systems (subgraph *A*), and (iii) competitive interactions between visual, semantic, ventral attention and subcortical systems (subgraph *O*). We found three subgraphs (*K, J, P*) with significantly greater expression bias for the Stroop task than expected by chance (*p* < 0.025; Bonferroni corrected for multiple comparisons). Specifically, subgraphs with significantly biased expression towards the Stroop task conditions exhibited: (i) cooperative interactions between auditory, subcortical and default mode systems (subgraph *K*), (ii) cooperative interactions between visual, cingulo-opercular, cerebellar, and dorsal attention systems (subgraph *J*), and (iii) competitive interactions between visual, fronto-parietal, dorsal attention, cingulo-opercular, and default mode systems (subgraph *P*).

To summarize, our results suggest that the Navon task exhibits a greater likelihood of recruiting cooperatively expressed subgraphs that overlap with executive control, visual and task-negative cognitive systems, and competitively expressed subgraphs that overlap with visual and somatosensory systems. In contrast, the Stroop task exhibits a greater likelihood of recruiting cooperatively expressed subgraphs that overlap with attention, salience, visual and task-negative cognitive systems, and competitively expressed subgraphs that overlap with executive control, visual, attention, salience and task-negative cognitive systems. Importantly, we note that in both the Stroop and Navon tasks the functional subgraphs expressed complex topological patterns of functional interactions that were distributed across several cognitive systems. Most notably, we identify subgraphs that suggest cooperation between regions in classically identified cognitive control and default mode regions during the Navon task and competition between these regions during the Stroop task. Critically, ten of the sixteen subgraphs were not significantly more expressed in any particular task than expected by chance and may implicate functional network components that are recruited for network processes that are agnostic to task-specific mechanics, such as arousal or cognitive control.

### 2.3. Subgraph expression adapts to transitions in cognitive demand

We next asked “How are functional subgraphs recruited in response to experimentally imposed changes in cognitive demand during the different cognitive control tasks?” We expected that subgraphs would be recruited differently during state transitions related to task initiation (as measured by the transition between fixation and the low cognitive demand condition) and during state transitions related to cognitive control (as measured by the transition between the low cognitive demand condition and the high cognitive demand condition) for the Stroop task and the Navon task. Specifically, we hypothesized that the transition in cognitive state that is mediated by cognitive control would be associated with coordinated shifts between cooperatively expressed subgraphs and competitively expressed subgraphs.

To test this hypothesis, we computed the relative change in each subgraph’s cooperative and competitive expression between the cognitive demand conditions of each task and averaged each subgraph’s relative measure across subjects. Intuitively, for transitions between the fixation condition and the low demand condition, a negative value implied greater expression of a subgraph during the fixation condition and a positive value implied greater expression of a subgraph during the low demand condition. Similarly, for transitions between the low demand condition and the high demand condition, a negative value implied greater expression of a subgraph during the low demand condition and a positive value implied greater expression of a subgraph during the high demand condition. To evaluate whether subgraphs predictably shift between cooperative and competitive expression during the transition between task conditions, we used a Spearman’s rank-order correlation test to compare the relative change in cooperative subgraph expression to the relative change in competitive subgraph expression during the transition between task conditions of each task (Fig. 5).

**Figure 5:**
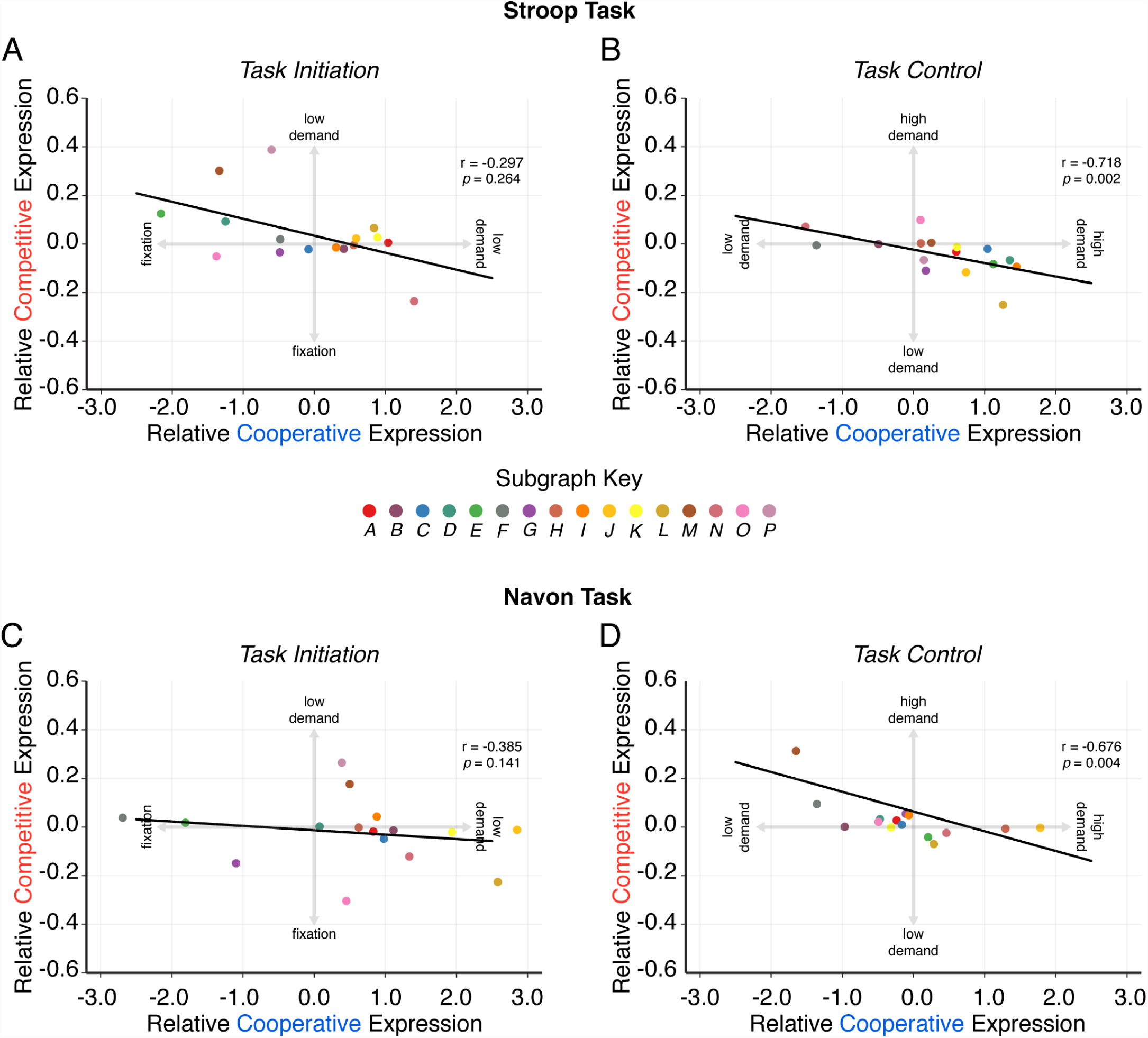
Antagonistic control strategy coincides with increased cognitive demand. Phase space of subgraph expression for task-specific transitions in cognitive demand. (*A*) Relative change in subgraph expression between the fixation condition and the low demand condition of the Stroop task for cooperative interactions (*x*-axis) and competitive interactions (*y*-axis), averaged over subjects. Negative values indicate greater expression during the fixation condition and positive values indicate greater expression during the low demand condition. Using a Spearman’s rank-order correlation test, we assessed whether a change in cognitive demand associated with Stroop task initiation predicts a change in the subgraphs’ functional roles between the fixation condition and the low demand condition. We found no significant relationship between the change in subgraph expression for cooperative interactions and the change in subgraph expression for competitive interactions (*ρ* = −0.30, *p* = 0.26). (*B*) We similarly compared the relative change in subgraph expression between the low demand condition and the high demand condition of the Stroop task for cooperative interactions and competitive interactions. Our results demonstrated that a change in cognitive demand associated with experimentally modulated task control in the Stroop task predicts a change in the subgraphs’ functional roles between the low demand condition and the high demand condition (*ρ* = −0.72, *p* = 0.002). These findings suggest that subgraphs flip between cooperative and competitive modes of expression as the Stroop task becomes more cognitively demanding. (*C*) For the Navon task, we found no significant relationship between the change in subgraph expression for cooperative interactions and the change in subgraph expression for competitive interactions during the transition to task initiation (*ρ* = −0.39, *p* = 0.14). (*D*) In contrast, we did find a significant relationship between the change in subgraph expression for cooperative interactions and the change in subgraph expression for competitive interactions during the transition to task control (*ρ* = −0.68, *p* = 0.004). These findings suggest that subgraphs flip between cooperative and competitive modes of expression as the Navon task becomes more cognitively demanding. Critically, our observation that the switch between cooperative and competitive modes of expression is evident only during task control conditions, implicate an adaptive network mechanism that is unique to cognitive control and not task initiation.

For the Stroop task, we found: (i) no significant relationship between cooperative and competitive relative subgraph expression during transitions between the fixation condition and the low demand condition (Fig. 5A; *ρ* = −0.30, *p* = 0.26; Bonferroni corrected for multiple comparisons), and (ii) a significant relationship between cooperative and competitive relative subgraph expression during transitions between the low demand condition and the high demand condition (Fig. 5B; *ρ* = −0.72, *p* = 0.002; Bonferroni corrected). These results suggest that shifts between cooperative and competitive subgraph expression accompany Stroop task transitions related to cognitive control but not task initiation. Similarly for the Navon task, we found: (i) no significant relationship between cooperative and competitive relative subgraph expression during transitions between the fixation condition and the low demand condition (Fig. 5C; *ρ* = −0.39, *p* = 0.14; Bonferroni corrected), and (ii) a significant relationship between cooperative and competitive relative subgraph expression during transitions between the low demand condition and the high demand condition (Fig. 5D; *ρ* = −0.68, *p* = 0.004; Bonferroni corrected). These results suggest that shifts between cooperative and competitive subgraph expression also accompany Navon task transitions related to cognitive control but not task initiation.

We next scrutinized the subgraphs that exhibited the greatest shifts in cooperative and competitive expression. During the Stroop task, we observed the greatest shift from cooperative to competitive involvement of the subgraph linking the frontoparietal, cingulo-opercular, and default mode systems (subgraph *N*) and the greatest shift from competitive to cooperative involvement of the subgraph linking the cingulo-opercular and fronto-parietal systems to visual, dorsal attention, and cerebellar systems (subgraph *L*). During the Navon task, we observed the greatest shift from cooperative to competitive involvement of the subgraph linking the cingulo-opercular system to the dorsal attention system (subgraph *M*) and the greatest shift from competitive to cooperative involvement of the subgraph linking dorsal attention, cerebellar, and visual systems (subgraph *J*). Together, these results suggest that traditional cognitive control, attentional, and cerebellar systems are most likely to shift between cooperative and competitive roles during the transition towards more cognitively demanding brain states.

Critically, this analysis demonstrated that functional subgraphs express coordinated changes in cooperative and competitive interaction during transitions to more cognitively demanding task conditions. Specifically, subgraphs that are more cooperatively expressed during low demand conditions become more competitively expressed during high demand conditions and subgraphs that are more competitively expressed during low demand conditions become more cooperatively expressed during high demand conditions. For both the Stroop task and the Navon task, we found that the functional brain network may utilize a control strategy that is marked by coordinating antagonistic changes in the co-activation between different cognitive systems.

### 2.4. Recruitment of functional subgraphs related to task performance

We next examined how recruitment of functional subgraphs relates to inter-individual performance during task initiation and during cognitive control. To evaluate an individual’s task performance, we separately computed each individual’s median reaction time over all low cognitive demand task trials and their median reaction time over all high cognitive demand task trials after regressing away the effect of motion. Intuitively, a lower reaction time here indicates better task performance. We interpret the reaction time on the low demand condition as a marker of task initiation, and we interpret the difference in reaction time between the high demand condition and the low demand condition as a marker of cognitive control. We next used the Spearman’s rank-order correlation test to study the relationship between relative change in a subgraph’s cooperative or competitive expression and individual performance related to task initiation and cognitive control (Fig. 6). To determine the statistical significance of each Spearman correlation test, we generated a null distribution of the correlation test statistic by randomly permuting the subgraph expression coefficients for the particular task conditions across all subgraph assignments and subjects. For each of 1000 permutations, we computed the relative subgraph expression between the task conditions, and then calculated the Spearman correlation between the surrogate relative subgraph expression values and individual task performance. We show a summary of the results from the statistical tests for the Stroop task in Table S2 and for the Navon task in Table S3.

**Figure 6:**
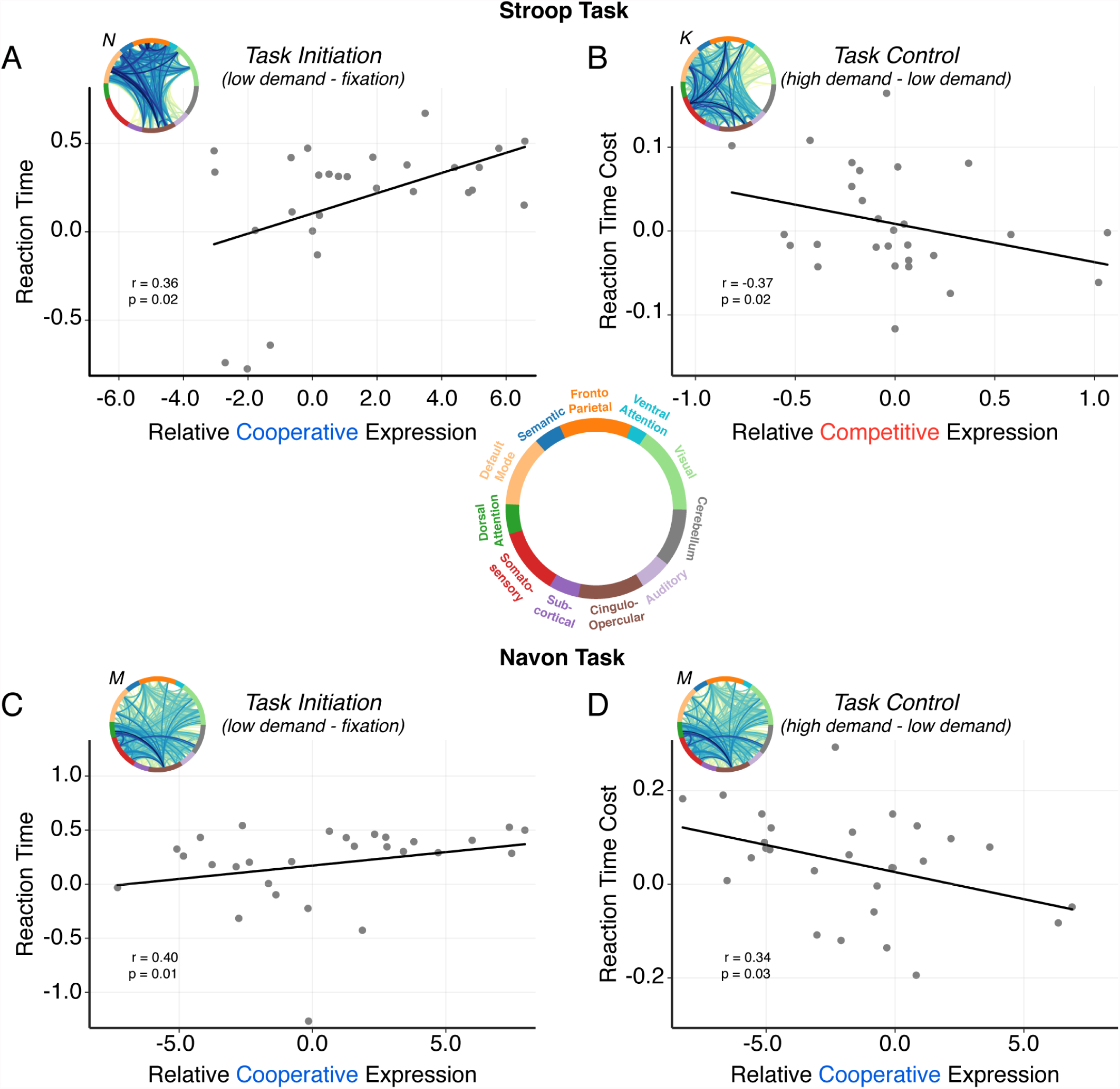
Subgraph expression stratifies inter-individual task performance. Relationship between the relative change in subgraph expression between task-specific, cognitive demand conditions and individual change in performance. (*A*) Increase in cooperative expression of subgraph *N* from the fixation condition to the low demand condition of the Stroop task is associated with increased reaction time on the low demand condition of the Stroop task (*ρ* = 0.36, *p* = 0.02), suggesting that subgraph *N* plays a negative role in behavioral performance during Stroop task initiation. (*B*) Increase in competitive expression of subgraph *K* from the low demand condition to the high demand condition of the Stroop task is associated with decreased reaction time cost from the low demand condition to the high demand condition of the Stroop task (*ρ* = 0.36, *p* = 0.02), suggesting that subgraph *K* plays a positive role in behavioral performance during Stroop task control. (*C*) Increase in cooperative expression of subgraph *M* from the fixation condition to the low demand condition of the Navon task is associated with increased reaction time on the low demand condition of the Navon task (*ρ* = 0.40, *p* = 0.01), suggesting that subgraph *M* plays a negative role in behavioral performance during Navon task initiation. (*D*) Increase in cooperative expression of subgraph *M* from the low demand condition to the high demand condition of the Navon task is associated with decreased reaction time cost from the low demand condition to the high demand condition of the Navon task (*ρ* = −0.34, *p* = 0.03), suggesting that subgraph *M* plays a positive role in behavioral performance during Navon task control.

In particular, for the Stroop task we found: (i) a significant positive relationship between the change in cooperative expression of subgraph *M* between the low demand condition and the fixation condition, and the reaction time on the low demand condition (Fig. 6A; *ρ* = 0.36, *p* = 0.02; uncorrected for multiple comparisons), and (ii) a significant negative relationship between the change in cooperative expression of subgraph *M* between the high demand condition and the low demand condition, and the change in reaction time between the high demand condition and the low demand condition (Fig. 6B; *ρ* = −0.37, *p* = 0.02). Intuitively, these results demonstrate that increased relative cooperative expression of subgraph *M* leads to poorer performance on the low demand condition of the Stroop task – suggesting that expression of this subgraph plays a negative role in behavioral performance during task initiation. Functionally, subgraph *N* is characterized by interactions between visual, cingulo-opercular, fronto-parietal, and default mode systems. Our results also demonstrate that increased relative competitive expression of subgraph *K* leads to better change in performance between the high demand condition and low demand condition – suggesting that expression of this subgraph plays a positive role in behavioral performance during cognitive control. Functionally, subgraph *M* is characterized by interactions between auditory, subcortical and default mode systems.

Similarly for the Navon task, we found: (i) a significant positive relationship between the change in cooperative expression of subgraph *M* between the low demand condition and the fixation condition, and the reaction time on the low demand condition (Fig. 6C; *ρ* = 0.40, *p* = 0.01), and (ii) a significant negative relationship between the change in competitive expression of subgraph *M* between the high demand condition and the low demand condition, and the change in reaction time between the high demand condition and the low demand condition (Fig. 6D; *ρ* = −0.34, *p* = 0.03). These results demonstrate that increased relative cooperative expression of subgraph *M* leads to poorer performance on the low demand condition of the Navon task – suggesting that expression of this subgraph plays a negative role in behavioral performance during task initiation. Conversely, increased relative cooperative expression of the same subgraph *M* lead to better change in performance between the high demand condition and low demand condition – suggesting that expression of this subgraph plays a positive role in behavioral performance during cognitive control. Functionally, subgraph *M* is characterized by interactions between cingulo-opercular, dorsal attention, somatosensory and cerebellar systems.

Lastly, we asked whether there are individual brain regions that are more likely to participate in subgraphs that facilitate or impede performance. We hypothesized that brain regions commonly associated with executive control, attention, and salience are more likely to behave as hubs in functional subgraphs that mediate task performance. To test this hypothesis, we first computed the node strength – a measure of the “hubness” of a node – by calculating the sum of all edge weights from a node for each of the 262 brain regions in each of the sixteen subgraphs. We next separately weighted the node strengths of each subgraph by their correlation values between relative expression and performance – that is nodes of the same subgraph were weighted by the same correlation value – for performance-positive subgraphs and performance-negative subgraphs. Finally, we computed the weighted sum of the node strengths for each brain region across subgraphs resulting in a performance-positive subgraph participation score and a performance-negative subgraph participation score for each brain region and each task condition. Intuitively, a greater participation score implies that a particular brain region is more influential in subgraphs associated with performance on a particular task condition. To determine whether a brain region exhibited a greater participation score than expected by chance, we constructed null distributions of regional participation scores for performance-positive and performance-negative subgraphs and each task condition by uniformly permuting the edges of each subgraph 1000 times and recomputing the regional participation score for each permutation. We retained regional participation scores that exceeded the 95% confidence interval of the null distribution after using Bonferroni correction for multiple comparisons testing (see Fig. S4 for real and null distributions of regional participation scores).

We then examined performance-positive and performance-negative regional participation scores for the task control conditions (transition from low demand to high demand) of the Stroop and Navon tasks (Fig. 7). For the Stroop task, we found that brain regions including lateral occipital cortex, superior parietal cortex, fusiform area, and rostral middle frontal gyrus were among the strongest hubs in performance-positive subgraphs, and we found that brain regions including precuneus, posterior cingulate cortex, paracingulate cortex, paracentral cortex, cuneus, and anterior cingulate cortex were among the strongest hubs in performance-negative subgraphs. These results indicate that during cognitive control conditions of the Stroop task, increased participation of regions involved in complex visual processing in their respective subgraphs may be more beneficial for performance and increased participation of regions involved in task salience may be more disadvantageous for performance. Similarly, for the Navon task we found that brain regions including cerebellum, rostral and caudal middle frontal gyri, and superior parietal cortex were among the strongest hubs in performance-positive subgraphs, and we found that brain regions including superior and inferior frontal gyri, inferior parietal cortex, and middle temporal gyrus were among the strongest hubs in performance-negative subgraphs. These results indicate that during cognitive control conditions of the Navon task, increased participation of regions involved in complex executive and sensorimotor processing in their respective subgraphs may be more beneficial for performance and increased participation of regions involved in awareness and executive functions may be more disadvantageous for perfor-mance.

**Figure 7:**
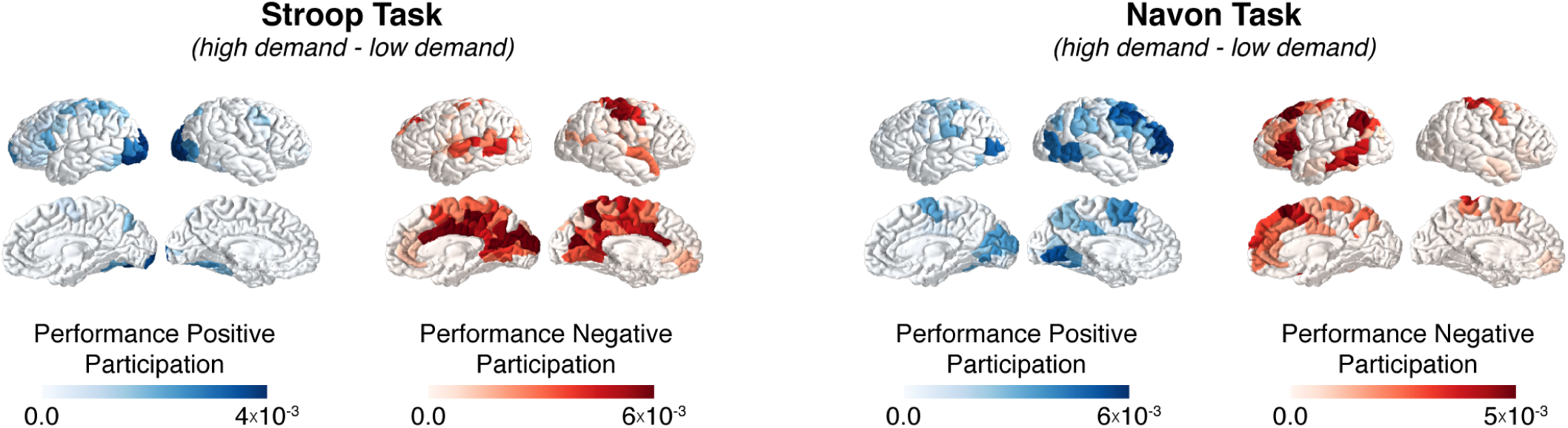
Regional participation in subgraph-mediated differences in task performance. Participation scores for individual brain regions that are significantly strong nodes in subgraphs that facilitate or impede task performance. To compute performance-positive and performance-negative participation scores, we computed the node strength of each brain region in each subgraph and calculated the sum of each brain region’s node strength across subgraphs, weighted by Pearson correlation value between subgraph expression and task performance. Intuitively, a greater participation score implies that a particular brain region is more influential in subgraphs associated with performance on a particular task condition. Participation scores were compared to a null distribution generated by permuting edges in the subgraph adjacency matrix 1000 times and recomputing the participation score for each permutation (*p* < 0.05; Bonferroni corrected for multiple comparisons). For the Stroop task, we found that brain regions including lateral occipital cortex, superior parietal cortex, fusiform area, and rostral middle frontal gyrus were among the strongest hubs in performance-positive subgraphs, and we found that brain regions including precuneus, posterior cingulate cortex, paracingulate cortex, paracentral cortex, cuneus, and anterior cingulate cortex were among the strongest hubs in performance-negative subgraphs. For the Navon task, we found that brain regions including cerebellum, rostral and caudal middle frontal gyri, and superior parietal cortex were among the strongest hubs in performance-positive subgraphs, and we found that brain regions including superior and inferior frontal gyri, inferior parietal cortex, and middle temporal gyrus were among the strongest hubs in performance-negative subgraphs. These results suggest that individual brain regions that are highly central in a subgraph’s topological architecture largely influence the relationship between the expression of the subgraph and task performance.

Together, these results demonstrate that objectively defined functional subgraphs that link distributed brain regions explain individual differences in performance related to task initiation and cognitive control in Stroop and Navon tasks. Importantly, our results further demonstrate that subgraphs may play either a facilitating or impeding role in behavioral performance during different phases of cognitive control tasks. Intriguingly, we also identified a potential non-linear constraint on Navon task performance whereby decreased expression of subgraph *M* supports task initiation but hinders effective cognitive control. This result elucidates an adaptive strategy for recruiting subgraphs during cognitive control whereby the functional goal associated with a set of network interactions can shift depending on cognitive state.

## 3. Discussion

In this work, we asked, “What functional constraints shape internally-guided transitions in brain state during cognitive control?” To answer this question, we applied a powerful machine-learning approach referred to as non-negative matrix factorization, to dynamic functional brain networks measured during two cognitive control tasks – yielding subgraphs or clusters of temporally co-varying functional interactions between brain regions. We studied the cooperative and competitive expression of these functional subgraphs as subjects transitioned between different levels of task-induced cognitive demand. We showed that the subgraphs differentiate clusters of functional interactions that are specific to the mechanics of the cognitive control tasks from those that are generalized to the network processes common to the cognitive control tasks. Specifically, we show for the first time clear evidence that functional subgraphs adaptively alter their cooperative and competitive expression depending on the type of cognitive control task and the amount of cognitive demand imposed on the system. Our results significantly extend our understanding of how objectively-defined clusters of functional interactions, beyond individual region-region co-activation, facilitate transitions between cognitive states.

### 3.1. Encoding dynamical rules for cognitive control

Our non-negative matrix factorization approach enabled us to objectively account for: (i) the dissociability of brain networks into composite subgraphs that are recruited for specific cognitive control functions, and (ii) the flexible and adaptive expression of these putative cognitive sub-networks in response to fluctuations in cognitive demand. Intuitively, these subgraphs represent clusters of functional interactions whose weights tend to fluctuate together across tasks and across conditions. Unlike other graph partitioning techniques, such as community detection, that pursue a hard partitioning of network nodes into discrete clusters, NMF enables a soft partitioning of the high dimensional set of network edges into subgraphs that allow an edge to participate in multiple network sub-units (Khambhati et al., 2016b; Chai et al., 2017). This capability is advantageous for examining how pairs of brain areas functionally interact within different topological contexts. Mathematically, NMF recovers a non-orthogonal basis set of graph edges whose linear combination – weighted by dynamic expression coefficients – can reconstruct the original space of observed network topologies across the experimental task conditions. In other words, subgraphs represent a set of functional relationships for the cognitive control data from which they were recovered and subgraph expression coefficients represent the encoding of those relationships for the different task conditions (we refer the reader to Olshausen and Field (1996) for a discussion on neural coding theory).

Thus from the perspective of network-based encoding of cognitive control tasks, indeed, we found that subgraphs are comprised of functional interactions that are either sensitive to the specific needs of a particular task or generalized to needs common over tasks. These data support the theory that there exists separate task-specific and task-general network architectures (Cole et al., 2014). We examined the particular cognitive systems involved in task-specific and task-general subgraphs and found a dual-role for cooperative and competitive interactions between traditional cognitive control systems and the default mode system – these systems are cooperatively expressed during cognitive control involving Navon-based task-shifting and are competitively expressed during cognitive control involving Stroop-based response inhibition. Our finding of competitive interactions between cognitive control and default mode systems during the Stroop task is well supported by the popular theory that the task-negative, default mode system deactivates as task-positive, executive areas activate (Raichle et al., 2001; Greicius et al., 2003; Fox et al., 2006). On the other hand, our finding of cooperative interactions between these systems during the Navon task challenges the notion that these systems must decouple during cognitive control. Prior studies have in fact demonstrated that individuals that exhibit greater cooperation between the default mode network and executive areas tend to display better behavioral performance during cognitive control tasks that involve switching between task-rules (Fornito et al., 2012; Crittenden et al., 2015). Based on these results, we posit that differences in the nature of functional interactions between these systems might be explained by task-specific requirements for cognitive control. Importantly, NMF demonstrated the ability to tease apart the functional interactions underlying intrinsic differences in cognitive control processes by recovering task-specific subgraphs.

### 3.2. Antagonistic push-pull control of cognitive demand

A growing body of literature in network neuroscience has shown that the brain possesses an ability to maintain a homeostasis of its own internal dynamics through antagonistic, pushpull interactions in various areas of healthy cognition (Fornito et al., 2012; Cocchi et al., 2013; Pezzulo et al., 2015) and disease (Khambhati et al., 2016a). Simply, push-pull control strategies may prevent imbalances of activity in complex, interconnected systems like the brain (Graybiel, 1996; He et al., 2014). A push-pull mechanism would be a critical component of cognitive control in which brain networks must perform two antagonistic functions: (i) segregated information processing in functionally-specific domains, and (ii) integrated information processing to adapt to environmentally-driven changes in cognitive demand (Shine et al., 2016). We sought to identify a potential push-pull control mechanism underlying the brain’s ability to shift between low and high cognitively demanding states by leveraging our ability to compare dynamical changes in cooperative and competitive expression of functional subgraphs. Our results demonstrated a “see-saw effect” as cognitive demand shifts between low and high demand states whereby: (i) subgraphs with more integrated and cooperative expression become more segregated and competitively expressed, and (ii) subgraphs with more segregated and competitive expression become more integrated and cooperatively expressed. We posit that a push-pull mechanism might internally regulate the extent of integration and segregation of different brain networks as they adaptively reorganize to meet cognitive demand. In our analysis, we observed that the transition between cognitively demanding brain states involves a change in the interacting roles between brain areas distributed across several cognitive systems: including executive, attentional, and cerebellar regions. Recent studies focusing on functional interactions between cerebellum and traditional cognitive control regions (Krienen and Buckner, 2009) have suggested that the cerebellum may sub-serve cognitive processes related to error correction (Dosenbach et al., 2008; Egner and Hirsch, 2005). Our results add new insight to this discussion by demonstrating in two different cognitive control tasks that: (i) cerebellum, executive, and sensory areas interact more cooperatively with increasing cognitive demand, and (ii) executive areas interact more competitively amongst themselves with increasing cognitive demand. In the context of push-pull control, these findings suggest that segregated information processing in executive areas is met with integrated information processing across distributed brain areas across executive, sensory and cerebellar systems.

We also considered the possibility that regulatory mechanisms involved in cognitive control might also explain differences in individual performance on cognitively demanding tasks. We found that subgraphs may be heterogeneously associated with individual cognitive performance – they may improve, impede, or minimally impact individual performance during both simple task conditions and demanding task conditions. These data suggest that cognitive control requires coordinated increases in the expression of particular clusters of functional interactions and coordinated decreases in the expression of other clusters to effectively facilitate task performance. In addition to regulating the degree of expression across different groups of subgraphs, other constraints may exist on the range of achievable expression levels of individual subgraphs. Specifically, we found that increased cooperation between executive and cerebellar areas was associated with better performance on a simple task condition and worse performance on a more cognitively demanding condition. Such an “inverted-U” shape performance curve has been previously cited as evidence for a constraint on the performance capacity that functional brain regions are capable of supporting (Callicott et al., 1999). We speculate that the rich distribution of performance-modes exhibited by functional subgraphs implicates a network homeostasis on cognitive control processes (Pezzulo et al., 2015).

### 3.3. Conclusions and future directions

In sum, we demonstrated that functional brain networks capably adapt their topological architecture in response to task-driven modulation in cognitive demand. Critically, we observed that cognitive control may not necessarily activate discrete cognitive brain systems, but rather recruit several interconnected systems, in concert, to facilitate transitions between cognitively demanding brain states. When individuals under- or over-express functional interactions between these cognitive systems they tend to respond more slowly during difficult cognitive tasks, implicating specific brain sub-networks in facilitating or impeding an individual’s ability to transition between states.

In this study, we focused on the mechanistic role that functional brain networks play in regulating internal dynamics during cognitive control. However, our novel approach and findings open new doors for querying how such regulatory mechanisms could be modulated to influence behavior. For instance, can we perturb specific network components to improve the likelihood that an individual is able to access shorter trajectories to switch between low demanding states and high demanding states? By marrying machine-learning approaches that objectively tease apart concurrent network processes attributed to different facets of cognition with burgeoning neurotechnologies such as neurofeedback (Bassett and Khambhati, 2017), neurostimulation (Cocchi et al., 2015), or pharmacological intervention (Schwarz et al., 2007; Smucny et al., 2014; Braun et al., 2016) that can exogenously control network dynamics, we can explore how disrupting network components that exhibit task-based adaptation causally influence behavior. The prospect of such scientific inquiry is equally exciting in diseases such as schizophrenia in which patients experience more probable transitions to more disruptive cognitive states.

## 4. Experimental Procedures

### 4.1. Study cohort

#### 4.1.1. Ethics Statement

All subjects volunteered with informed consent in writing in accordance with the Institutional Review Board/Human Subjects Committee at the University of Pennsylvania.

#### 4.1.2. Patient Demographics

A total of 30 subjects were recruited. All subjects were screened for prior history of psychiatric or neurological illness. One subject was excluded due to near-chance performance on the task (accuracy = 52%). One additional subject was excluded due to technical problems on the day of scanning. The final sample included 28 individuals (mean age = 25.6 ± 3.5, 70% caucasian, 13 females).

### 4.2. Cognitive Control Tasks

All participants completed a Stroop task with color-word pairings that were eligible and ineligible to elicit interference effects (Stroop, 1935) and a local-global perception task based on classical Navon figures (Navon, 1977). For the Stroop task, trials were comprised of words presented one at a time at the center of the screen printed in one of four colors – red, green, yellow, or blue - on a gray background. For all trials, subjects responded using their right hand with a four-button response box. All subjects were trained on the task outside the scanner until proficient at reporting responses using a fixed mapping between the color and button presses (i.e., index finger = “red”, middle finger = “green”, ring finger = “yellow”, pinky finger = “blue”). Trials were presented in randomly intermixed blocks containing trials that were either eligible or ineligible to produce color-word interference effects. In the scanner, blocks were administered with 20 trials apiece separated by 20 s fixation periods with a black crosshair at the center of the screen. Each trial was presented for a fixed duration of 1900 ms separated by an interstimulus interval of 100 ms during which a gray screen was presented. In the trials ineligible for interference, the words were selected to not conflict with printed colors (“far,” “horse,” “deal,” and “plenty”). In the trials eligible for interference (i.e. those designed to elicit the classic Stroop effect (Stroop, 1935)), the words were selected to introduce conflict (i.e., printed words were “red,” “green,” “yellow,” and “blue” and always printed in an incongruent color). In our analysis, we refer to blocks that are eligible (ineligible) to produce color-word interference effects as *high demand* (*low demand*) conditions (**Fig. 1B**).

For the Navon task, local-global stimuli were comprised of four shapes – a circle, X, triangle, or square – that were used to build the global and local aspects of the stimuli. On all trials, the local feature did not match the global feature, ensuring that subjects could not use information about one scale to infer information about another. Stimuli were presented on a black background in a block design with three blocks. In the first block type, subjects viewed white local-global stimuli. In the second block type, subjects viewed green local-global stimuli. In the third block type, stimuli switched between white and green across trials uniformly at random with the constraint that 70% of trials included a switch in each block. In all blocks, subjects were instructed to report only the local features of the stimuli if the stimulus was white, and to report only the global feature of the stimuli if the stimulus was green. Blocks were administered in a random order. Subjects responded using their right hand with a four-button response box. All subjects were trained on the task outside the scanner until proficient at reporting responses using a fixed mapping between the shape and the button presses (i.e., index finger = “circle”, middle finger = “X”, ring finger = “triangle”, pinky finger = “square”). In the scanner, blocks were administered with 20 trials apiece separated by 20 s fixation periods with a white crosshair at the center of the screen. Each trial was presented for a fixed duration of 1900 ms separated by an interstimulus interval of 100 ms during which a black screen was presented. In our analysis, we refer to blocks that switch between local-global perception as the *high demand* condition and blocks that do not switch as the *low demand* condition (Fig. 1C).

### 4.3. Data Acquisition and Pre-Processing

We acquired T1-weighted anatomical scans on a Siemens 3.0T Tim Trio for all subjects. Anatomical scans were segmented using FreeSurfer (Fischl, 2012) and parcellated using the connectome mapping toolkit (Cammoun et al., 2012) into *N* = 234 cortical and subcortical brain regions. We also included a cerebellar parcellation (*N* = 28 brain regions Diedrichsen et al. (2009)) by using FSL to nonlinearly register the individuals T1 to MNI space. Then, we used the inverse warp parameters to warp the cerebellum atlas to the individual T1. Finally, we merged the cerebellar label image with the dilated cortical and subcortical parcellation image resulting in *N* = 262 brain regions.

Functional magnetic resonance imaging was acquired on a 3.0T Siemens Tim Trio whole-body scanner with a whole-head elliptical coil by means of a single-shot gradient-echo T2* (TR = 1500 ms; TE = 30 ms; flip angle = 60 degrees; FOV = 19.2 cm, resolution 3mm x 3mm x 3mm). Preprocessing was performed using FEAT v. 6.0 (fMRI Expert Analysis Tool) a component of the FSL software package (Jenkinson et al., 2012). To prepare the functional images for analyses, we completed the following steps: skull-stripping with BET to remove non-brain material, motion correction with MCFLIRT (FMRIBs Linear Image Registration Tool; (Jenkinson et al., 2012)), slice timing correction (interleaved), spatial smoothing with a 6-mm 3D Gaussian kernel, and high pass temporal filtering to reduce low frequency artifacts. We also performed EPI unwarping with fieldmaps to improve subject registration to standard space. Native image transformation to a standard template was completed using FSL’s affine registration tool, FLIRT (Jenkinson et al., 2012). Subject-specific functional images were co-registered to their corresponding high-resolution anatomical images via a Boundary Based Registration technique (BBR (Greve and Fischl, 2009)) and were then registered to the standard MNI-152 structural template via a 12-parameter linear transformation. Finally, each participant’s individual anatomical image was segmented into grey matter, white matter, and CSF using the binary segmentation function of FAST v. 4.0 (FMRIBs Automated Segmentation Tool (Zhang et al., 2001)). The white matter and CSF masks for each participant were then transformed to native functional space and the average timeseries were extracted. Images were spatially smoothed using a kernel with a full-width at half-maximum of 6 mm. These values were used as confound regressors on the time series along with 18 translation and rotation parameters as estimated by MCFLIRT (Jenkinson et al., 2002).

We refer the reader to Medaglia et al. (2016b) for additional methodological details regarding data acquisition and pre-processing.

### 4.4. Constructing Functional Brain Networks

We constructed functional brain networks to study the functional interactions between brain regions during the Stroop and Navon cognitive control tasks. To measure functional interactions, we first separately divided the BOLD signal into six low demand blocks, six high demand blocks, and twelve fixation blocks (before each cognitive demand block) for each behavioral task of each subject – each block contained 20 samples or 30 seconds of signals (Fig. 2B). We next computed a Pearson correlation similarity measure between each pair of BOLD signals from the *N* brain regions (graph nodes) in each of the *K* experimental blocks and aggregated correlations (graph edges) into an *N* × *N* × *K* adjacency matrix **A** for each subject.

To analyze cooperative (positively correlated) and competitive (negatively correlated) functional interactions, we separated positively-weighted edges from negatively-weighted edges for each block *k* in **A**. This procedure resulted in a thresholded adjacency matrix **A**\* of size *N* × *N* × 2 × *K* where each block *k* is associated with one *N* × *N* matrix with positive edge weights and another *N* × *N* matrix with negative edge weights (Fig. 2C).

An alternate representation of the adjacency matrix **A**\* is a two-dimensional network configuration matrix **Â**\*, which tabulates all *N* × *N* pairwise edge weights across *K* blocks, and across cooperative and competitive edge types (**Fig. 2D**). Due to symmetry of **A**\*_*k*_, we unravel the upper triangle of **A**\*_*k*_, resulting in the weights of *N*(*N* − 1)*/*2 connections. Thus, **Â**\* has dimensions *N*(*N* − 1)/2 × 2 * *K*.

### 4.5. Clustering Functional Networks into Subgraphs

To identify network subgraphs – sets of network edges whose strengths co-vary over experimental task conditions – we applied an unsupervised machine learning algorithm called non-negative matrix factorization (NMF) (Lee et al., 1999) to the network configuration matrix. This technique enabled us to pursue a parts-based decomposition of the network configuration matrix into subgraphs with expression coefficients that vary with time (Fig. 2E,F). For in-depth discussion regarding network subgraphs, we refer the reader to (Khambhati et al., 2016b). For recent applications of subgraph decomposition to the study of functional brain networks, please see (Chai et al., 2017; Khambhati et al., 2017). Specifically, we first computed the magnitude of the network configuration matrix **Â**\* such that all entries of the matrix were non-negative. We next solved the matrix factorization problem **Â**\* ≈ **WH**s.t.**W** > = 0, **H** > = 0 by optimizing the following cost function:

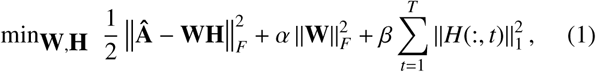

where *m* ∊ [2, min(*N*(*N* − 1)/2, *T*) − 1] is the number of subgraphs to decompose, *β* is a penalty weight to impose sparse temporal expression coefficients, and *α* is a regularization of the interaction strengths for subgraphs (Kim and Park, 2011). To solve the NMF equation, we used an alternating non-negative least squares with block-pivoting method with 100 iterations for fast and efficient factorization of large matrices (Kim et al., 2014). We initialized **W** and **H** with non-negative weights drawn from a uniform random distribution on the interval [0, 1].

To select the parameters *m*, *β*, and *α*, we pursued a random sampling scheme – shown to be effective in optimizing high-dimensional parameter spaces (Bergstra and Bengio, 2012; Khambhati et al., 2016b) – in which we re-ran the NMF algorithm for 1000 parameter sets in which *m* is drawn from 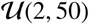, *β* is drawn from 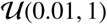, and *α* is drawn from 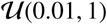 (Fig. 2). We evaluated subgraph learning performance based on a fourteen-fold cross-validation scheme in which the twenty eight subjects are uniformly partitioned into folds of two subjects and, iteratively, thirteen folds are used to identify subgraphs and the held-out fold is used to compute the cross-validation error (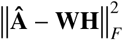). The optimal parameter set should yield subgraphs that minimize the cross-validation error and reliably span the space of observed network topologies (Khambhati et al., 2016b). Based on these criteria, we identified an optimum parameter set (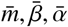) that exhibited a low residual error in the bottom 25*^th^* percentile of our random sampling scheme (Fig. 2G-I).

Due to the non-deterministic nature of this approach, we integrated subgraph estimates over multiple runs of the algorithm using *consensus clustering* – a general method of testing robustness and stability of clusters over many runs of one or more non-deterministic clustering algorithms (Monti et al., 2003). Our adapted consensus clustering procedure entailed the following steps: (i) run the NMF algorithm *R* times per network configuration matrix, (ii) concatenate subgraph matrix **W** across *R* runs into an aggregate matrix with dimensions 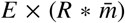, and (iii) apply NMF to the aggregate matrix to determine a final set of subgraphs **W**_consensus_ and expression coefficients **H**_consensus_ (we refer the reader to (Khambhati et al., 2016b) for more details). In this study, we set *R* = 100.

## Supplementary Tables and Figures

**Figure 1:**
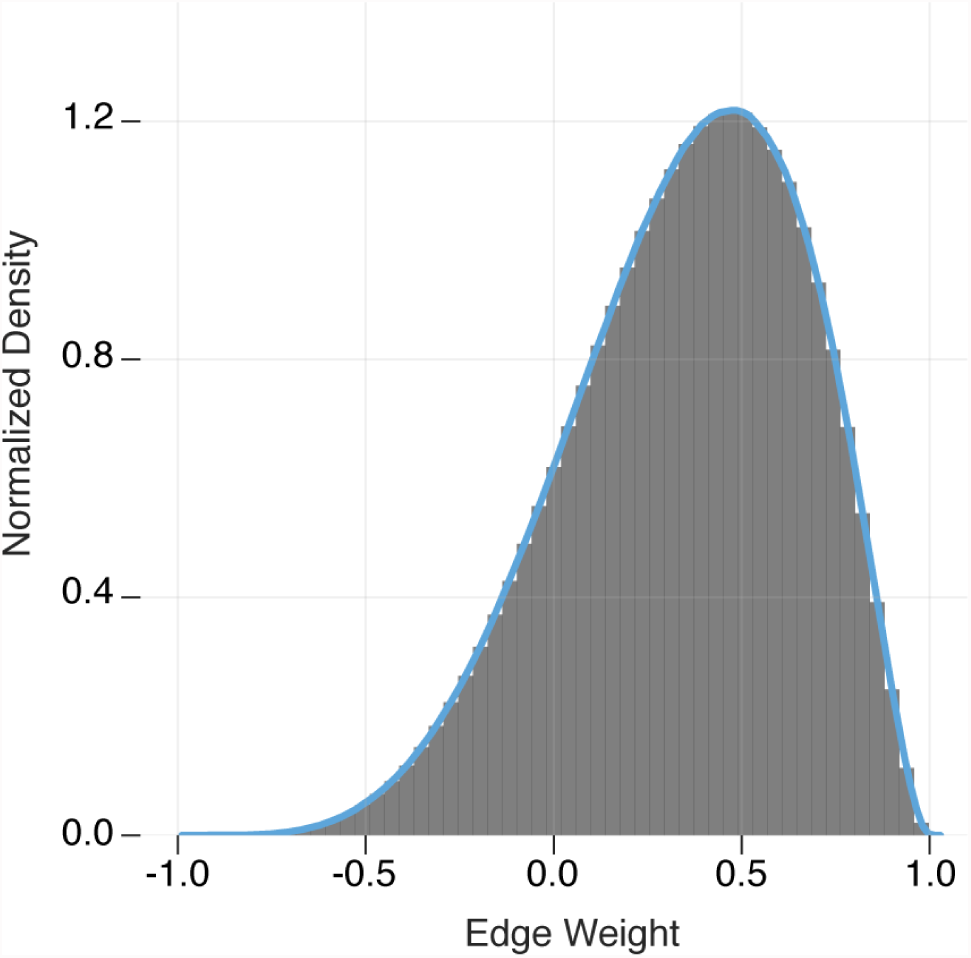
Distribution of edge weights in the dynamic functional network. Dynamic functional networks were measured using a Pearson correlation similarity statistic between the BOLD fMRI time-series of 262 brain regions. To ensure negative correlation values were not introduced artifactually, we did not regress the global mean signal from the BOLD fMRI signal of each brain region [5]. We visualized the distribution of edge weights across all subjects and time windows to confirm that the measured functional connectivity indeed exhibited both positive and negative values.

**Figure 2:**
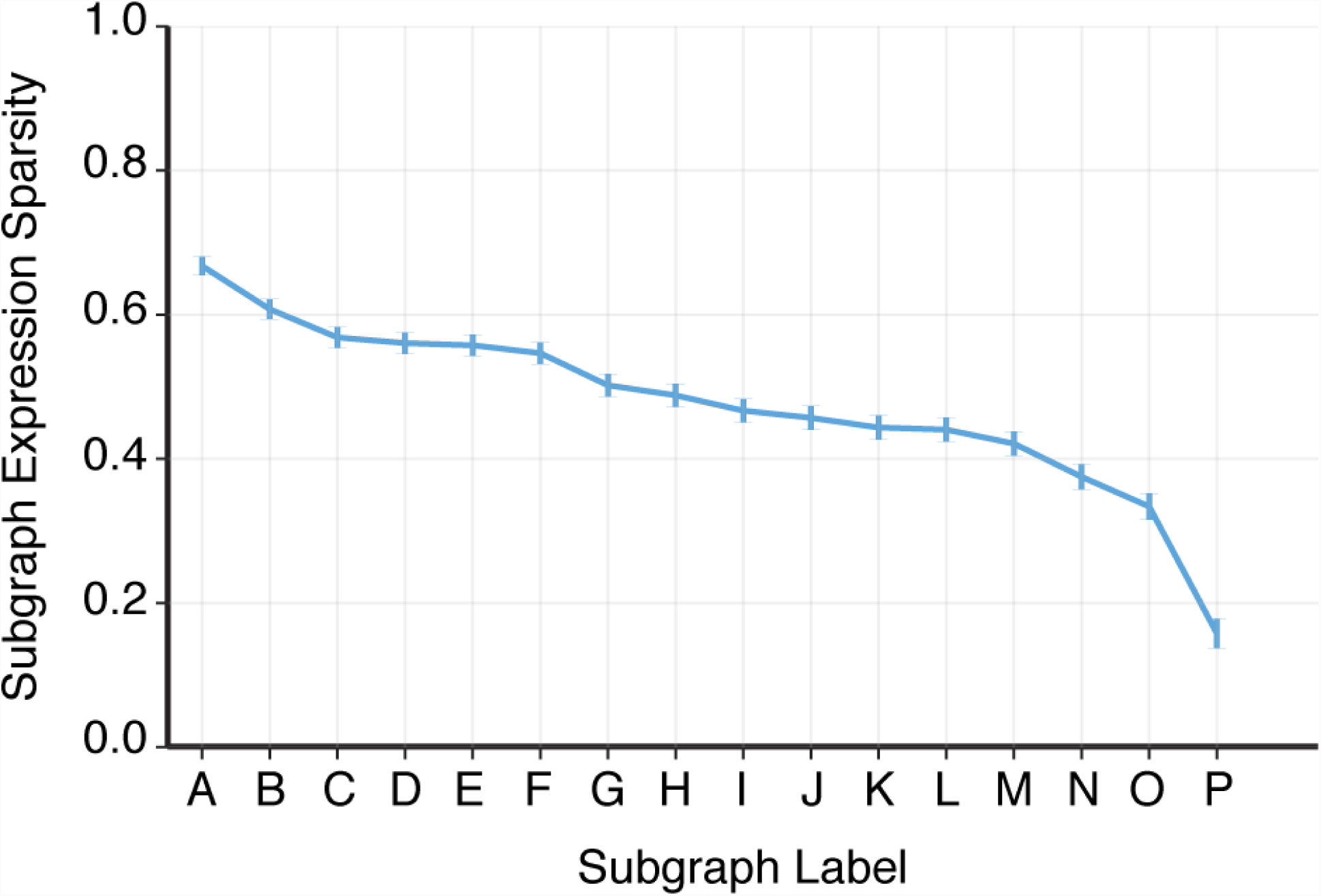
Ranking subgraphs based on sparsity of expression coefficients. Subgraphs were assigned a letter ranging from *A-P* depending on the sparsity of their expression coefficients. Specifically, we computed the mean number of experimental blocks (across tasks, task conditions, and subjects) with an expression weight of zero and ranked subgraphs in decreasing order. Intuitively, subgraphs with greater expression sparsity are more specific in their expression to particular experimental blocks, and subgraphs with less expression sparsity are more generalized in their expression over experimental blocks. We refer to subgraphs based on this letter assignment throughout our study. Error bars represent standard error over the mean.

**Figure 3:**
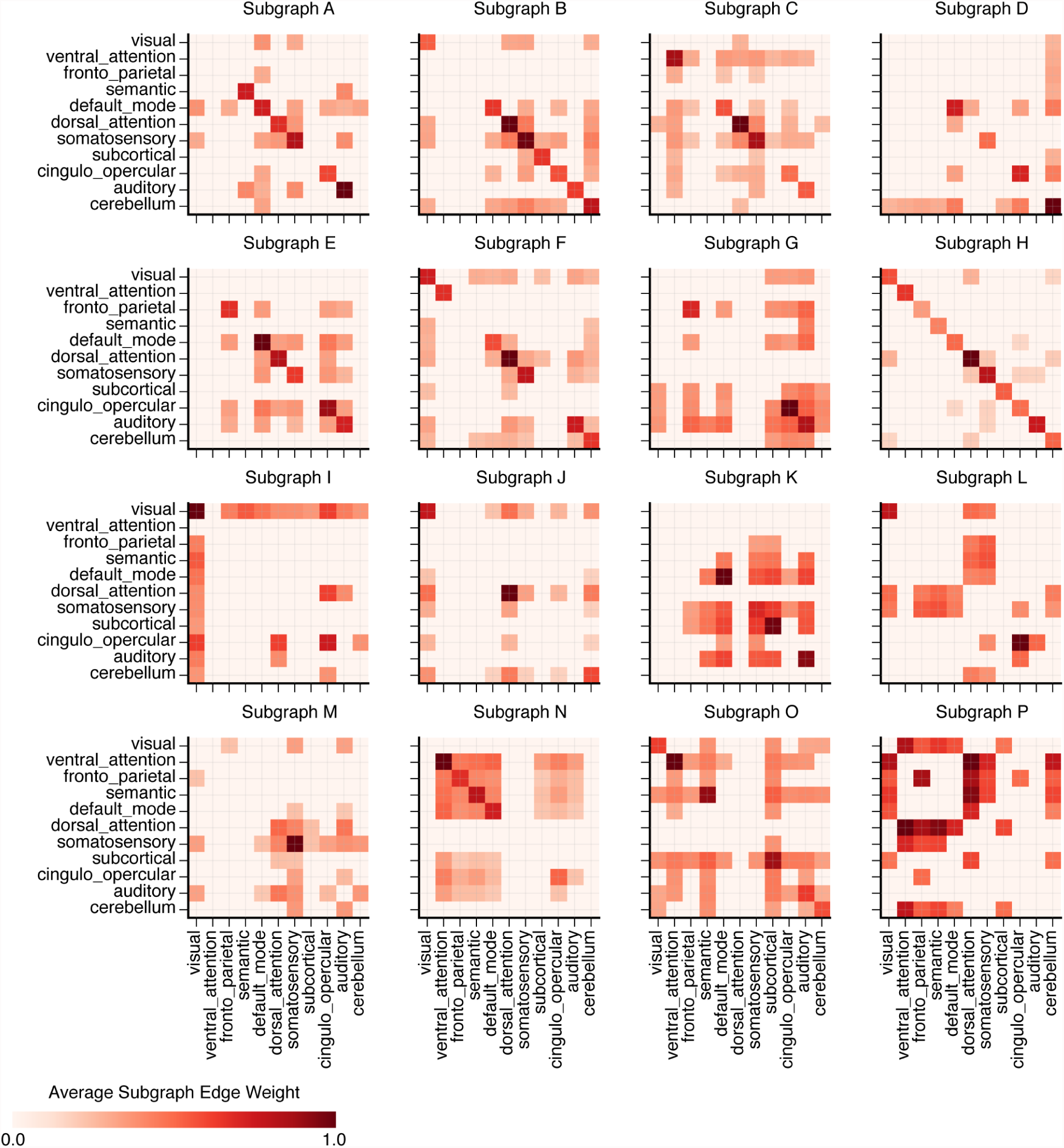
Functional subgraphs capture distributed interactions between cognitive systems. We determined whether functional subgraphs reflect functional interactions within and between known cognitive brain systems using a previously documented approach [2]. Briefly, this procedure enabled us to map each subgraph’s 262 × 262 regional-level adjacency matrix to an 11 × 11 systems-level adjacency matrix - 262 refers to the number of brain regions in the anatomical atlas and 11 refers to the following cognitive systems [4]: frontoparietal, cingulo-opercular, dorsal attention, ventral attention, default mode, somatosensory, auditory, visual, subcortical, cerebellum, and semantic (see Table S1 for specific region-to-system assignments). Specifically, for each subgraph we computed the average edge weight between nodes assigned to the same cognitive system – represented by the diagonal elements of the systems-level adjacency matrix – and the average edge weight between all possible pairs of nodes assigned to two different cognitive systems - represented by the off-diagonal elements of the systems-level adjacency matrix. To assess whether a within-system or between-system interaction was more likely observed due to the topology of the subgraph than expected by chance, we generated a null distribution for each system-level interaction for each subgraph by permuting a subgraph’s edge weights between nodes 1000 times and recomputing the average edge weight for each permutation. We then compared each true system-level interaction to the null distribution and retained only significant interactions (*p* < 0.05; Bonferroni corrected for multiple comparisons). As a result of this procedure, we observed that each subgraph exhibited multiple within- and between-system interactions that were more likely than expected 5 by chance.

**Figure 4:**
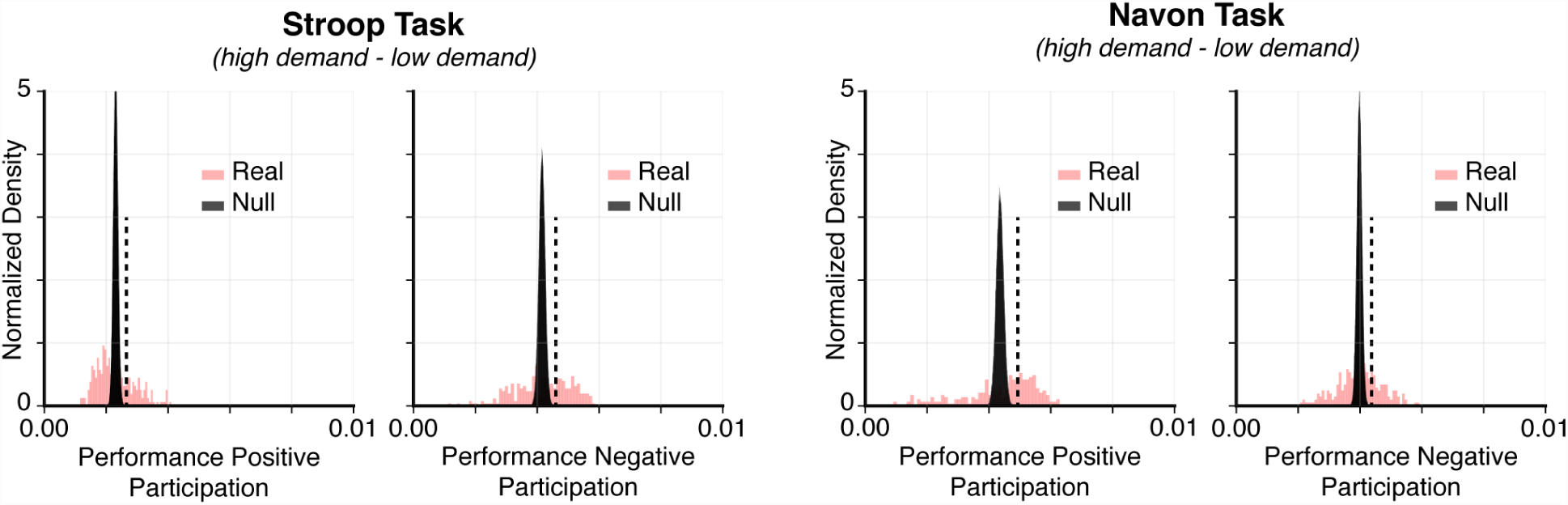
Identifying regional participation scores in functional subgraphs. To test whether particular brain regions are more likely to be centralized hubs in subgraphs that are associated with task performance, we measured the performance-weighted subgraph participation score for each brain region and task condition. Specifically, we first computed the node strength – the sum of all edge weights stemming from a node - for each brain region and each subgraph. We next separately weighted the node strengths of each subgraph by their correlation values between relative expression and performance – that is nodes of the same subgraph were weighted by the same correlation value – for performance-positive subgraphs and performance-negative subgraphs. We finally computed the weighted sum of the node strengths for each brain region across subgraphs resulting in a performance-positive subgraph participation score and a performance-negative subgraph participation score for each brain region and each task condition. Intuitively, a greater participation score implies that a particular brain region is more influential in subgraphs associated with performance on a particular task condition. To determine whether a brain region exhibited a greater participation score than expected by chance, we constructed null distributions of regional participation scores for performance-positive and performance-negative subgraphs and each task condition by uniformly permuting the edges of each subgraph 1000 times and recomputing the regional participation score for each permutation. We retained regional participation scores that exceeded the 95% confidence interval of the null distribution after using Bonferroni correction for multiple comparisons testing. Real distribution of participation scores are shown in red, and the null distribution of participation scores are shown in gray. Vertical lines demarcate the Bonferroni adjusted confidence interval.

**Table 1:** Assignments of individual brain regions to cognitive systems. We parcellated the brain into 262 regions of interest in which cortical and subcortical brain areas were delineated by [1] and cerebellar brain areas were delineated by [3]. Brain regions were assigned to cognitive brain systems based on a previously documented mapping by [4].

| Cognitive System | Parcellated Brain Region |
| --- | --- |
| Auditory | banks of the superior temporal sulcus<br>superior temporal<br>transverse temporal |
| Cerebellum | cerebellum |
| Cingulo Opercular | caudal anterior cingulate<br>rostral anterior cingulate<br>pars opercularis<br>pars orbitalis<br>rostral middle frontal<br>supramarginal |
| Default Mode | isthmus cingulate<br>posterior cingulate<br>precuneus<br>superior frontal |
| Dorsal Attention | superior parietal |
| Fronto Parietal | caudal middle frontal<br>frontal pole<br>inferior parietal<br>insula<br>medial orbitofrontal<br>pars triangularis<br>medial orbitofrontal<br>pars triangularis |
| Somatosensory | paracentral<br>postcentral<br>precentral |
| Subcortical | accumbens area<br>amygdala<br>caudate<br>hippocampus<br>globus pallidus<br>putamen<br>thalamus |
| Ventral Attention | lateral orbitofrontal |
| Visual | cuneus<br>entorhinal<br>fusiform<br>inferior temporal<br>lateral occipital<br>lingual<br>pericalcarine |
| Semantic | brainstem<br>middle temporal<br>parahippocampal<br>temporal pole |

**Table 2:** Relationship between subgraph expression and performance on the Stroop task. A Spearman’s rank-order correlation test was used to compare the relative change in subgraph expression during Stroop task conditions to the change in median reaction time over all trials of each task condition. Positive correlations imply increased subgraph expression was related to poorer performance, and negative correlations imply increased subgraph expression was related to better performance. Significant correlations highlighted in red were assessed at *p* < 0.05, uncorrected for multiple comparisons.

| Subgraph | Low Demand - Fixation<br>Cooperative Expression | Low Demand - Fixation<br>Competitive Expression | High Demand - Low Demand<br>Cooperative Expression | High Demand - Low Demand<br>Competitive Expression |
| --- | --- | --- | --- | --- |
| A | $\rho=0.19, p=0.15$ | $\rho=0.05, p=0.38$ | $\rho=0.21, p=0.14$ | $\rho=-0.05, p=0.39$ |
| B | $\rho=0.26, p=0.08$ | $\rho=0.29, p=0.06$ | $\rho=0.24, p=0.10$ | $\rho=0.12, p=0.26$ |
| C | $\rho=-0.08, p=0.34$ | $\rho=0.20, p=0.15$ | $\rho=0.01, p=0.47$ | $\rho=-0.37, p=0.02$ |
| D | $\rho=-0.16, p=0.20$ | $\rho=0.24, p=0.10$ | $\rho=-0.06, p=0.36$ | $\rho=-0.05, p=0.39$ |
| E | $\rho=0.18, p=0.17$ | $\rho=-0.07, p=0.36$ | $\rho=0.06, p=0.36$ | $\rho=-0.09, p=0.32$ |
| F | $\rho=-0.04, p=0.40$ | $\rho=0.00, p=0.48$ | $\rho=0.09, p=0.32$ | $\rho=-0.07, p=0.34$ |
| G | $\rho=-0.03, p=0.42$ | $\rho=0.10, p=0.30$ | $\rho=0.03, p=0.44$ | $\rho=-0.19, p=0.14$ |
| H | $\rho=-0.15, p=0.21$ | $\rho=-0.03, p=0.43$ | $\rho=-0.24, p=0.10$ | $\rho=0.10, p=0.28$ |
| I | $\rho=-0.18, p=0.17$ | $\rho=-0.01, p=0.47$ | $\rho=-0.19, p=0.16$ | $\rho=0.31, p=0.05$ |
| J | $\rho=0.01, p=0.46$ | $\rho=0.09, p=0.29$ | $\rho=-0.17, p=0.18$ | $\rho=-0.25, p=0.09$ |
| K | $\rho=0.02, p=0.45$ | $\rho=-0.15, p=0.21$ | $\rho=0.16, p=0.20$ | $\rho=-0.37, p=0.02$ |
| L | $\rho=-0.07, p=0.36$ | $\rho=0.27, p=0.07$ | $\rho=0.00, p=0.48$ | $\rho=0.12, p=0.26$ |
| M | $\rho=0.08, p=0.34$ | $\rho=-0.08, p=0.34$ | $\rho=0.00, p=0.48$ | $\rho=0.10, p=0.29$ |
| N | $\rho=0.36, p=0.02$ | $\rho=-0.00, p=0.47$ | $\rho=0.08, p=0.33$ | $\rho=-0.02, p=0.45$ |
| O | $\rho=0.05, p=0.37$ | $\rho=-0.08, p=0.32$ | $\rho=-0.05, p=0.39$ | $\rho=-0.04, p=0.41$ |
| P | $\rho=0.08, p=0.34$ | $\rho=0.09, p=0.31$ | $\rho=-0.19, p=0.15$ | $\rho=-0.00, p=0.50$ |

**Table 3:**
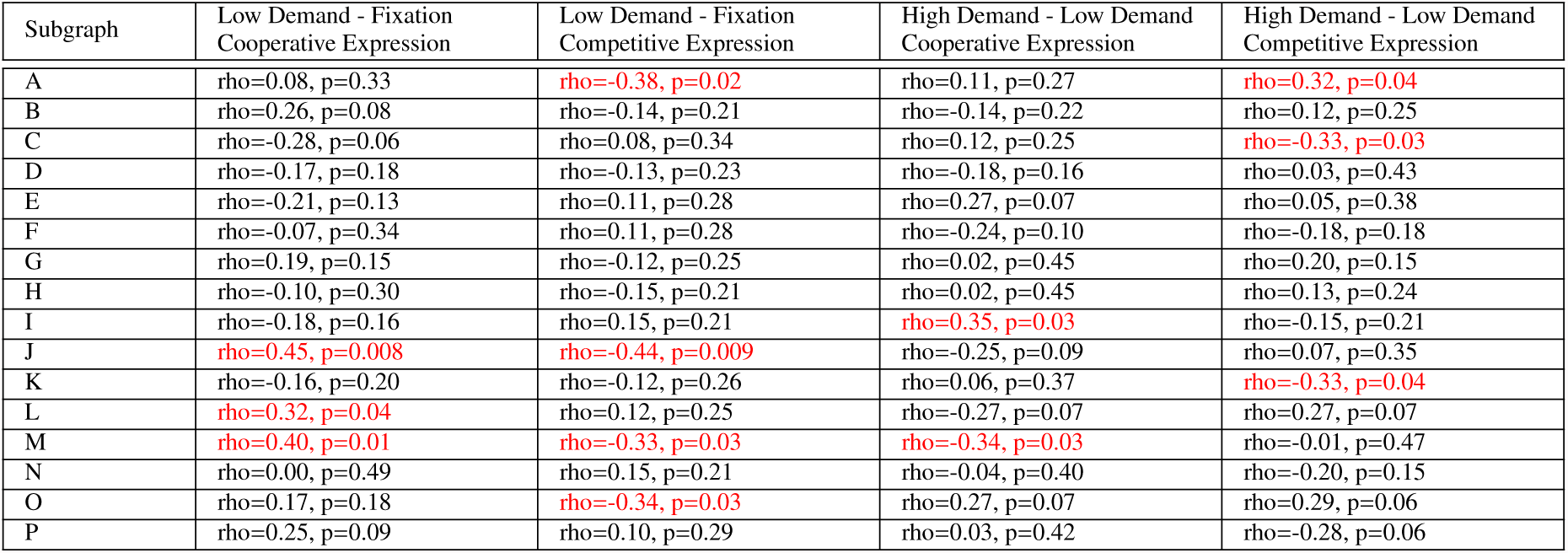
Relationship between subgraph expression and performance on the Navon task. A Spearman’s rank-order correlation test was used to compare the relative change in subgraph expression during Navon task conditions to the change in median reaction time over all trials of each task condition. Positive correlations imply increased subgraph expression was related to poorer performance, and negative correlations imply increased subgraph expression was related to better performance. Significant correlations highlighted in red were assessed at *p* < 0.05, uncorrected for multiple comparisons.

